# Generating an Executable Model of the Drosophila Central Complex

**DOI:** 10.1101/051318

**Authors:** Lev E. Givon, Aurel A. Lazar

## Abstract

The central complex (CX) is a set of neuropils in the center of the fly brain that have been implicated as playing an important role in vision-mediated behavior and integration of spatial information for locomotor control. In contrast to currently available data regarding the neural circuitry of neuropils in the fly's vision and olfactory systems, comparable data for the CX neuropils is relatively incomplete; many categories of neurons remain only partly characterized, and the synaptic connectivity between CX neurons has yet to be experimentally determined. Successful modeling of the information processing functions of the CX neuropils therefore requires a means of easily constructing and testing a range of hypotheses regarding both the high-level structure of their neural circuitry and the properties of their constituent neurons and synapses. This document demonstrates how NeuroArch and Neurokernel may be used to algorithmically construct and evaluate executable neural circuit models of the CX neuropils and their interconnects based upon currently available information regarding the geometry and polarity of the arborizations of identified local and projection neurons in the CX.

## I. Introduction

The brain of the fruit fly *Drosophila melanogaster* comprises approximately 50 neuropils. Most of these modules - referred to as local processing units (LPUs) are characterized by unique populations of local neurons; some - called hubs - do not contain any local neurons 3. The central complex (CX) comprises between 2,000 and 5,000 neurons [4] organized in four neuropils: the protocerebral bridge (PB), fan-shaped body (FB), ellipsoid body (EB), and noduli (NO) (Fig. 1). The former three neuropils contain local neurons, while the latter does not [3]. In contrast to most neuropils in the fly brain, PB, FB, and EB are unpaired; NO comprises either 3 [5] or 4 [4] paired neuropils. When considered as components of a single structure, FB and EB are together referred to as the central body (CB). Accessory brain areas that are connected directly to neuropils in CX include AL, BU, CRE, IB, LAL, SMP, WED, and PS [1, Fig. 7] (see § 2.1 for the neuropil nomenclature).

**Fig. 1.**
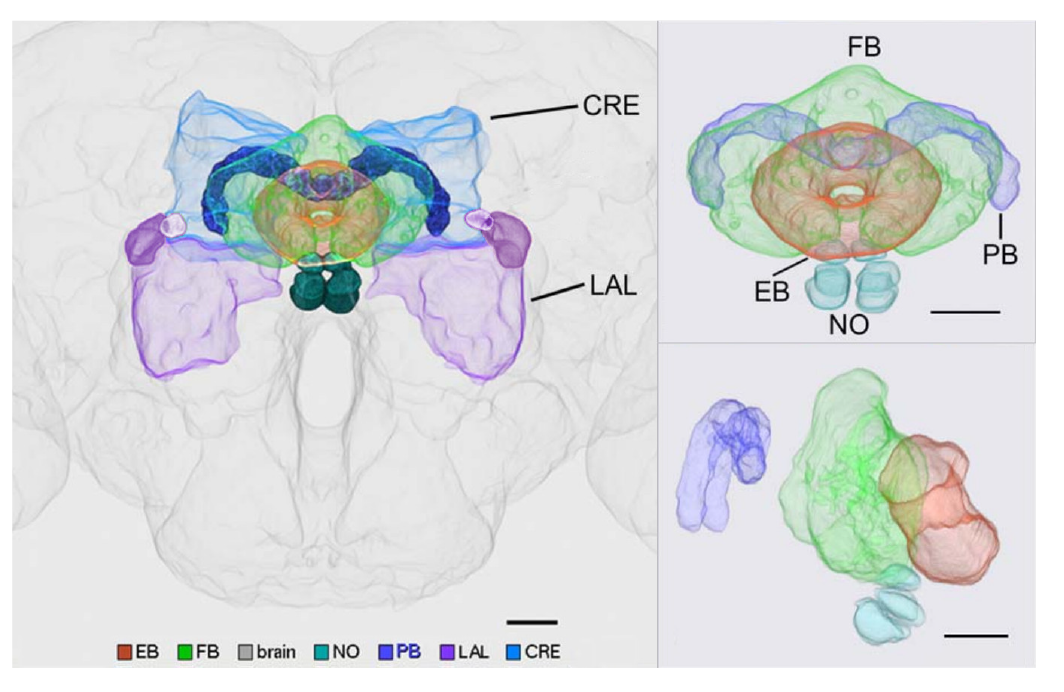
Volumetric image of central complex (PB, FB, EB, NO) and some accessory neuropils (CRE, LAL) arborized by CX neurons [5]. (©2015 Wiley Periodicals, Inc.)

Genetic experiments have shown that the CX neuropils play essential roles in a range of important behaviors:

i. EB appears to be involved in visual place learning [6], short-term orientation memory [7], [8], path integration, and left-right bargaining [4, p. 3];
ii. FB appears to also play a role in left-right bargaining, as well as visual pattern memory and object recognition [4, p. 8];
iii. PB plays a role in controlling step length and hence direction of walking [4, p. 7];
iv. NO neuropils seem to be involved in flight control [6, p. 179].

While some functional models of the CX neuropils have been presented [4], they do not explicitly show how the CX circuitry implements the information processing functions associated with the above behaviors or how the various neuropils’ individual functions combine to produce more comprehensive behaviors such as long-term motor skill learning or locomotor activity control.

Studies of neural activity in the CX of the fruit fly and arthropods have examined the response both of unidentified and specific CX neurons to visual inputs [9], [10], [11], [12]; owing to the limitations of electrophysiological techniques, these studies only examined the responses of individual neurons during each trial. Simultaneous observation of the aggregate neural responses of a set of isomorphic neurons in the fruit fly EB to visual input signals was achieved using calcium imaging [8], [13] (Fig. 2).

**Fig. 2.**
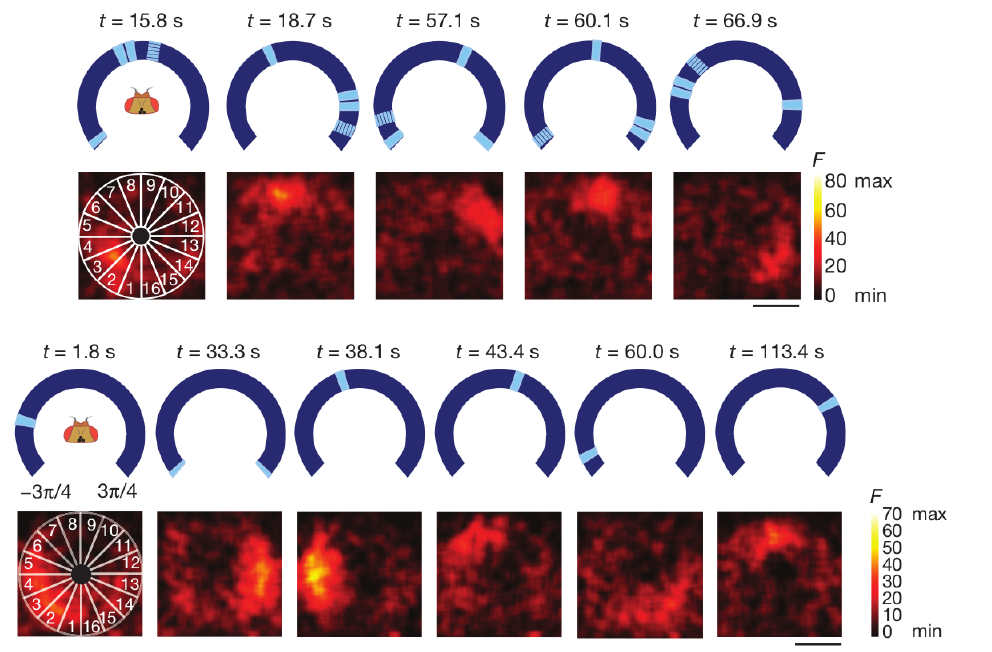
Observed activity of EB-LAL-PB neuron arborizations (§ 5.4.3) in EB in response to visual input at different points on the azimuth [13] (©2015 Macmillan Publishers Ltd.)

Connectome information for the CX neuropils is currently less complete than that available for sensory neuropils such as those in the fly olfactory and vision systems, the latter of which has recently been mapped in great detail using electron microscopy [14]. Although a range of local and projection neurons in CX have been identified and grouped into isomorphic sets using Golgi staining, genetic tagging techniques, and confocal microscopy [15, 16, 1, 5], many other CX neurons have not been systematically characterized and the synaptic connectivity between them remains unknown owing to the limitations of the above optical imaging technologies and the very limited EM-based analysis of CX synapses done to date (for an example of the latter, see [2]). This ambiguity regarding the structure of the CX neural circuitry compounds the already difficult task of modeling a portion of the brain that does not receive direct sensory input.

In light of the incompleteness of the CX connectome, it is perhaps unsurprising that only a few computational models of the CX neuropils or the entire CX currently exist. A spiking neural network model of spatial memory formation and storage in EB is presented in [17]; while this model can replicate experimental results for specific behaviors using a ring attractor circuit inspired by that of EB, it does not attempt to account for the exact observed biological circuitry or explain how such a model interacts with the other CX neuropils. A model of CX was included in a more comprehensive insect brain simulation described in [18], but it employs generalized models of the CX neuropils that use artificial behavior selection networks which - although they superficially make use of spiking neuron models - do not employ the observed neural circuitry of the neuropils.

To enable further investigation of the information processing capabilities of the CX neuropils, we need to be able to efficiently generate and evaluate a range of different executable CX models given the limited available connectome data. While a similar approach involving *C. elegans* has been used to generate an ensemble of testable models regarding the neural basis for salt klinotaxis behavior [19], the greater total number of neurons in the fruit fly CX and the need to evaluate the CX models together with models of the neuropils that provide them with input calls for

i. a database-driven approach to systematically creating an ensemble of CX different models of the neural circuitry that attempt to describe those characteristics of the circuitry that are as yet unknown, and
ii. a means of integrating models of these CX neuropil models with those of accessory neuropils from which they receive input into a single executable partial brain model.

This RFC is organized as follows. We first present the nomenclature used to identify neuropils and a scheme for labeling neurons in terms of their arborization patterns in § 2. We then review known data regarding the structure of neuropils in and connected to CX in § 3, and describe what is known regarding input pathways into the CX neuropils and the nature of their neural responses in § 4. We then present the arborization patterns of identified CX neurons in § 5. Finally, we describe a scheme for construction of an executable circuit model of CX that utilizes this arborization data and specified structural assumptions to infer the presence of synapses in § 6 and show how NeuroArch and Neurokernel expedite the process of executable circuit construction and evaluation.

## II. Terminology

### 2.1 Neuropil Nomenclature

*Drosophila* neuropils are identified in this document using the nomenclature described in [20]. Some neuropils are referred to by different names either in other literature or in other insects; [20, Tab. S13] maps the employed nomenclature to that used in [3] and - for the most part - in [1]. For neuropils that occur in pairs, upper case denotes the neuropil on the left side of the fly brain (from a dorsal perspective the fly) while lower case denotes the neuropil on the right side of the fly brain.

- Antennal Lobe (AL).
- Anterior optic tubercle (AOTU) - Also known as optic tubercle (OPTU [3]).
- Antler (ATL) - Corresponds to dorsal part of caudalcentral protocerebrum (CCP) [20, Tab. S13].
- Bulb (BU) - Also known as lateral triangle (LT, LAT, Lat Tri [1, Tab. S13] or LTR [15]).
- Crepine (CRE) - Posterior part also known as dorsal part of IDFP [20, Tab. S13]; comprises a region called the rubus (RUB) [5, p. 1001] or round body (RB) [5, p. 1031].
- Lateral accessory lobe (LAL) - Also known ventral body (VBO [15]) or inferior dorsofrontal protocerebrum (IDFP) [1, Fig. S1]. Comprises the gall (GA) [20, Tab. S13], whose dorsal and ventral portions are referred to as the dorsal and ventral spindle bodies (DSB and VSB, respectively) [5, p. 1021].
- Ellipsoid body (EB) - Also known as lower central body (CBL [6]).
- Fan-shaped body (FB) - Also known as upper central body (CBU [6]).
- Inferior Bridge (IB) - Corresponds to ventral part of caudalcentral protocerebrum (CCP) [20, Tab. S13].
- Lobula (LO).
- Lobula Plate (LOP).
- Noduli (NO).
- Posterior slope (PS) - Corresponds to caudalmedial protocerebrum (CMP) and - possibly - part of the ventromedial protocerebrum (VMP) [20, Tab. S13].
- Protocerebral Bridge (PB).
- Superior medial protocerebrum (SMP) - Corresponds to superior dorsolateral protocerebrum (SDFP) and medial part of inner dorsolateral protocerebrum (IDLP) [20, Tab. S13].
- Ventrolateral protocerebrum (VLP) - Contains optic glomeruli [20, p. 42, Supp.].
- Wedge (WED) - Also known as the caudal ventrolateral protocerebrum (CVLP) [20, p. 42, Supp.].

### 2.2 Neuron Labeling

Most neurons innervating the various CX and accessory neuropils possess at least two distinct clusters of dendrites (postsynaptic terminals) and/or axons (presynaptic terminals) that occupy geometrically distinct regions of the innervated neuropils [15]. These clusters are referred to as arborizations (Fig. 3). Since many CX neurons belong to distinct sets of morphologically similar neurons with similar arborization patterns, it is useful to use the latter to uniquely label each CX neuron type. If neurotransmitter profiles are ignored and each CX neuron type is assumed to be represented by a single neuron, then each neuron’s label unambiguously encodes the geometric regions of its arborizations and whether each arborization contains dendrites, axons, or both. This labeling scheme can be described in terms of the following parsing expression grammar (PEG) [21]; the grammar may be used to extract the arborizations of a particular neuron for constructing models of the CX circuitry (e.g., by using overlapping presynaptic and postsynaptic arborizations to infer synaptic connectivity). Note that a special case for handling the string LRB in the (name) rule (which corresponds to the left RB region of CRE) is necessary to prevent that string from being incorrectly parsed into LB and RB.

**Fig. 3.**
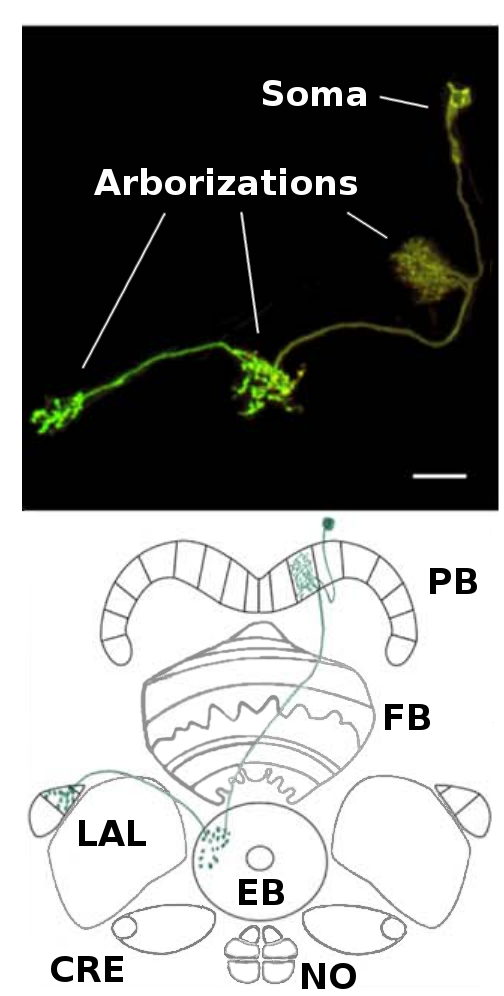
Example of neuron arborizations for a PB-EB-LAL neuron (§ 5.4.8) [5]. Each of the neuron arborizations occupies a specific region in different neuropils. (©2015 Wiley Periodicals, Inc.)

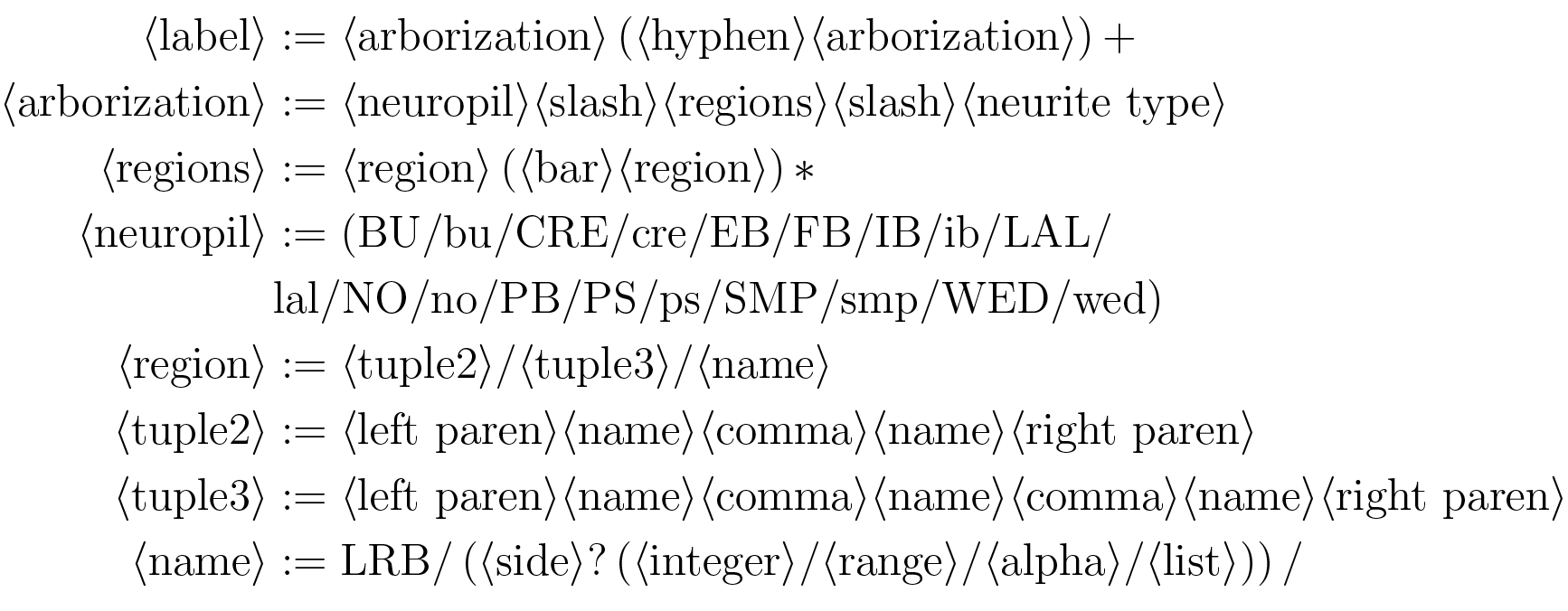

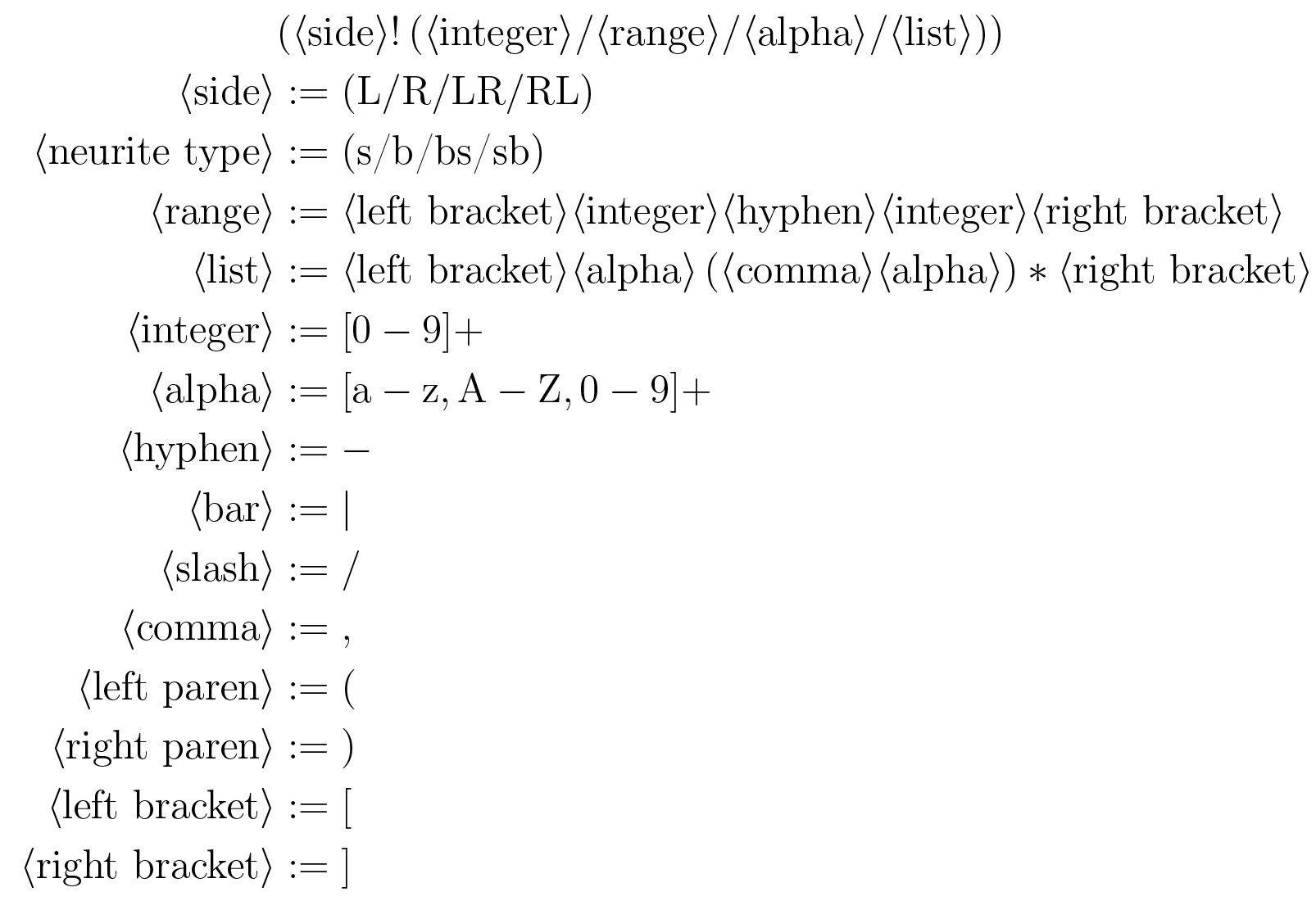

Neuropils are denoted by their abbreviated names as specified in § 2.1 and [20]; regions or compartments within neuropils are described and assigned names in § 3. The neurite type may be spine (s), bouton (or bleb) (b), or a combination thereof (bs, sb). In the absence of detailed data regarding synapses, information flow polarity is assumed to be reflected by neurite type; spines are assumed to be postsynaptic (and accept input), while boutons are assumed to be presynaptic (and emit output) [5, p. 1002]. Left and right are assumed to be with respect to a dorsal view of the fly.

## 3. Structure of Neuropils in and Associated with the Central Complex

This section presents details regarding the high-level structure of the various CX neuropils and the accessory neuropils to which they are connected.

### 3.1 Protocerebral Bridge (PB)

The PB neuropil comprises 18 regions called glomeruli [5] connected to other substructures within the CX (Fig. 4). The local neuron population of the PB comprises 8 [5, p. 1007] or 10 [1, p. 1743] types of local neurons (Fig. 12,Table 7). A single PB region label matches the following *regular expression*:

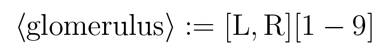

**Fig. 4.**
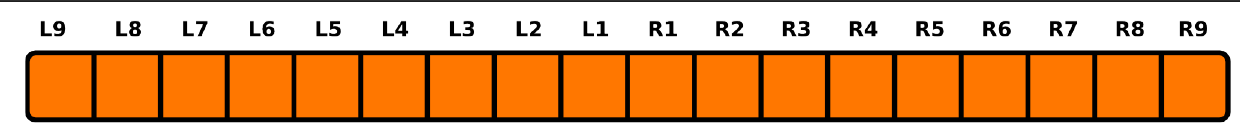
Schematic of regions in PB used to identify neurons by their arbors [1, 5].

### 3.2 Fan-Shaped Body (FB)

The FB neuropil comprises multiple lateral layers [6]; most recent work suggests the presence of 9 layers [5, p. 1011]. The neuropil is subdivided vertically into 8 [1] or 7 [5, p. 1010] columns called segments [15]; however, it seems that only some of its layers (1-5) exhibit clearly columnar structure [5, p. 1008] (5). Regions in FB are connected by local neurons called *pontine neurons;* some of these neurons connect adjacent layers, while others connect adjacent segments [15, p. 349, 352]. A representative class of pontine neurons comprising symmetric neurons that connect each segment in one side of FB with each segment in the other side such that the presynaptic and postsynaptic arborizations are 4 segments apart [15, p. 352] (although more recent work suggests that each neuron might be a bundle of 2 neurons [22, p. 1439]) is depicted in Tab. 8. Other classes dorsoventrally connecting different layers in FB may exist, but they have not been systematically identified. A single region in FB is denoted by a label matching the *regular expression*

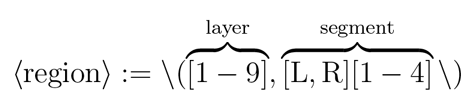

**Fig. 5.**
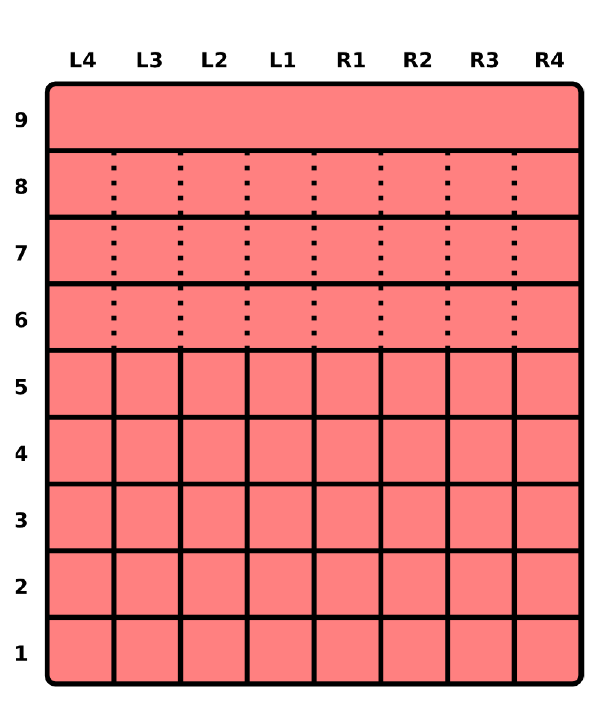
Schematic of regions in FB used to identify neurons by their arbors [1, 5].

### 3.3 Ellipsoid Body (EB)

The EB neuropil is a toroidal structure that comprises 16 wedges [5, p. 1013] (Fig. 6a), 8 tiles [5, p. 1018] (Fig. 6b), 3 shells (anterior, medial, posterior) [5, p. 1013], and 4 rings [1] (Fig. 6c). Wedges extend radially through full radius of the EB torus and occupy the posterior and medial shells or all 3 shells [5, p. 1013]. Tiles are restricted to the posteriorEllipsoid Body (EB) shell [5, p. 1014]; tiles geometrically overlap with corresponding wedges as described in Table 1. Although EB appears to contain local neurons [3], these neurons have not yet been systematically identified; there is some evidence for EB pontine neurons in related fly species such as *Neobellieria* [11, p. 11]. Each region in EB is denoted by a label that matches the *regular expression*

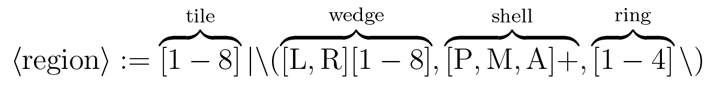

For EB regions other than tiles, the region denoted by a label comprises the volume intersected by the specified wedges, shells, and rings. For example, (L1, [P, M], 4) represents the volume in which wedge L1, shells P and M, and ring 4 overlap.

**Table I.** Geometric overlap between EB tiles and wedges.

| Tile | Wedge |
| --- | --- |
| EB/1/x | EB/([L1,R1],P,x)/x |
| EB/2/x | EB/(R[2,3],P,x)/x |
| EB/3/x | EB/(R[4,5],P,x)/x |
| EB/4/x | EB/(R[6,7],P,x)/x |
| EB/5/x | EB/([R8,L8],P,x)/x |
| EB/6/x | EB/(L[6,7],P,x)/x |
| EB/7/x | EB/(L[4,5],P,x)/x |
| EB/8/x | EB/(L[2,3],P,x)/x |

**Fig. 6.**
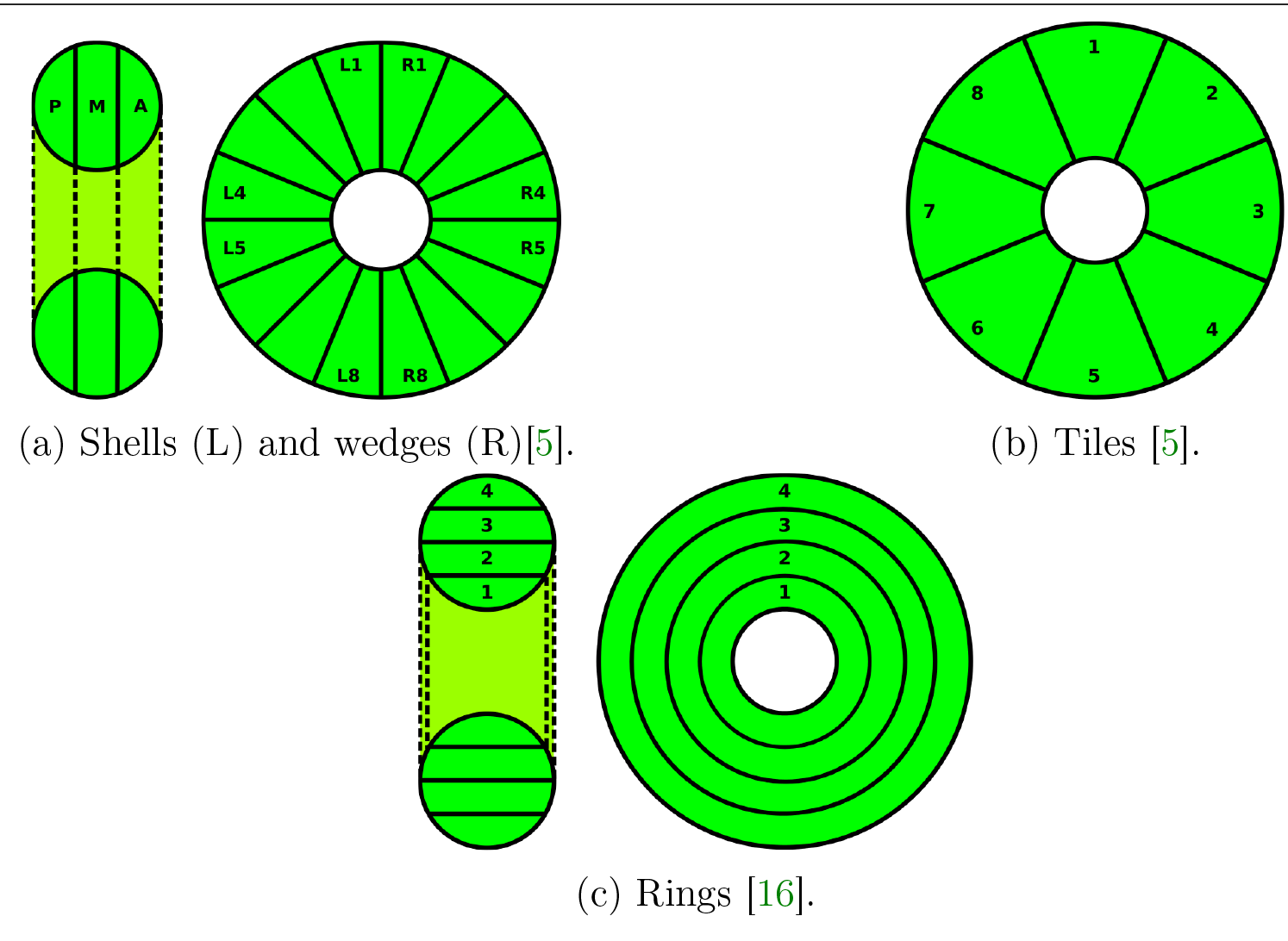
Schematics of regions in EB. All circular schematics are anterior; sagittal views in Fig. 6a and Fig. 6c are posterior to anterior from left to right).

### 3.4 Noduli (NO)

The NO neuropils comprise 3 distinct structures (N01, N02, N03) divided into subcompartments (Fig. 7) [5, p. 1017]. In contrast to the other CX neuropils, the noduli lack segregated populations of local neurons [3, p. 5]. Each N0 region label matches the following *regular expressions*:

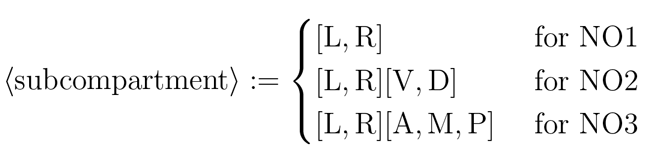

**Fig. 7.**
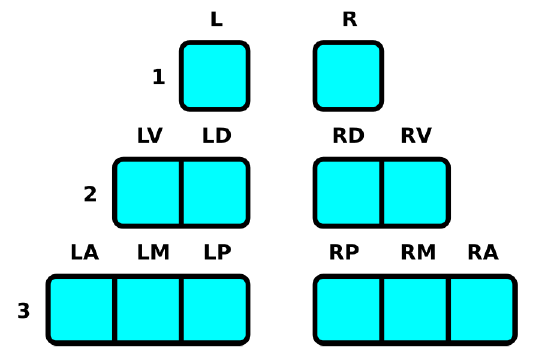
Schematic of regions in N0 used to identify neurons by their arbors [5].

### 3.5 Bulb (BU)

Each of the BU neuropils comprises multiple regions referred to as microglomeruli. There appear to be 80 microglomeruli in each BU neuropil (Fig. 8) [1, p. 1741]. These microglomeruli ostensibly exhibit retinotopic organization [8]. Each of the BU region labels matches the *regular expression*

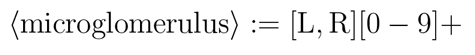
where the integer portion of the labels ranges from 1 to 80.

**Fig. 8.**
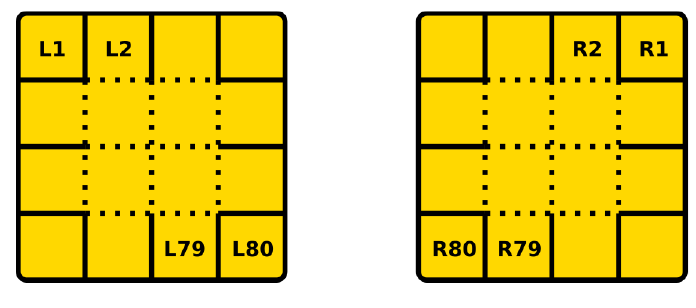
Schematic of regions in BU used to identify neurons by their arbors [1]. The relative positions of the regions does not necessarily correspond to their actual physical positions.

### 3.6 Lateral Accessory Lobe (LAL)

Each LAL neuropil comprises a region called the gall that is subdivided into a tip, dorsal, and ventral subregion; the remainder of LAL is referred to as the hammer body (HB) (Fig. 9) [1]. Each of these regions has a label that matches the *regular expression*

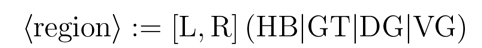

**Fig. 9.**
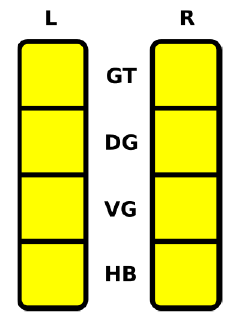
Schematic of regions in LAL used to identify neurons by their arbors [5].

### 3.7 Crepine (CRE)

Each CRE neuropil is divided into two regions (Fig. 10); these match the *regular expression*

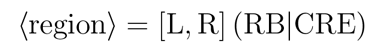

**Fig. 10.**
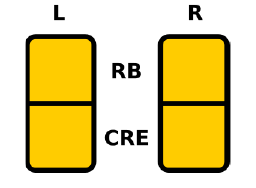
Schematic of regions in CRE used to identify neurons by their arbors [5].

### 3.8 Other Neuropils (IB, PS, SMP, WED)

Distinct regions of interest within IB, PS, SMP, and WED have not been identified; they are therefore regarded as comprising single regions on each side of the fly brain. Each region in these neuropils matches the *regular expression*

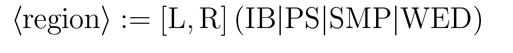

## IV. Central Complex Input Pathways and Neuron Responses

The neuropils in the CX are connected to various neuropils, but evidently not to any that directly receive sensory input except the antennal lobe (AL) [15, Fig. 24a]. Apart from connections between the CX neuropils and the accessory neuropils depicted in Fig. 11, connections have been observed between superior/inferior protocerebra and FB, between AOTU and BU [23, p. 9], and between VLP and PB [24, p. 9]. Preprocessed visual data from LO appears to enter the EB from BU via AOTU [25, p. 939], while additional visual input enters PB from other optic glomeruli in VLP [24, p. 9]. Other input enters FB via LAL. There also seems to be evidence of CX receiving mechanosensory information from the fly’s legs [4, p. ^6^].

**Fig. 11.**
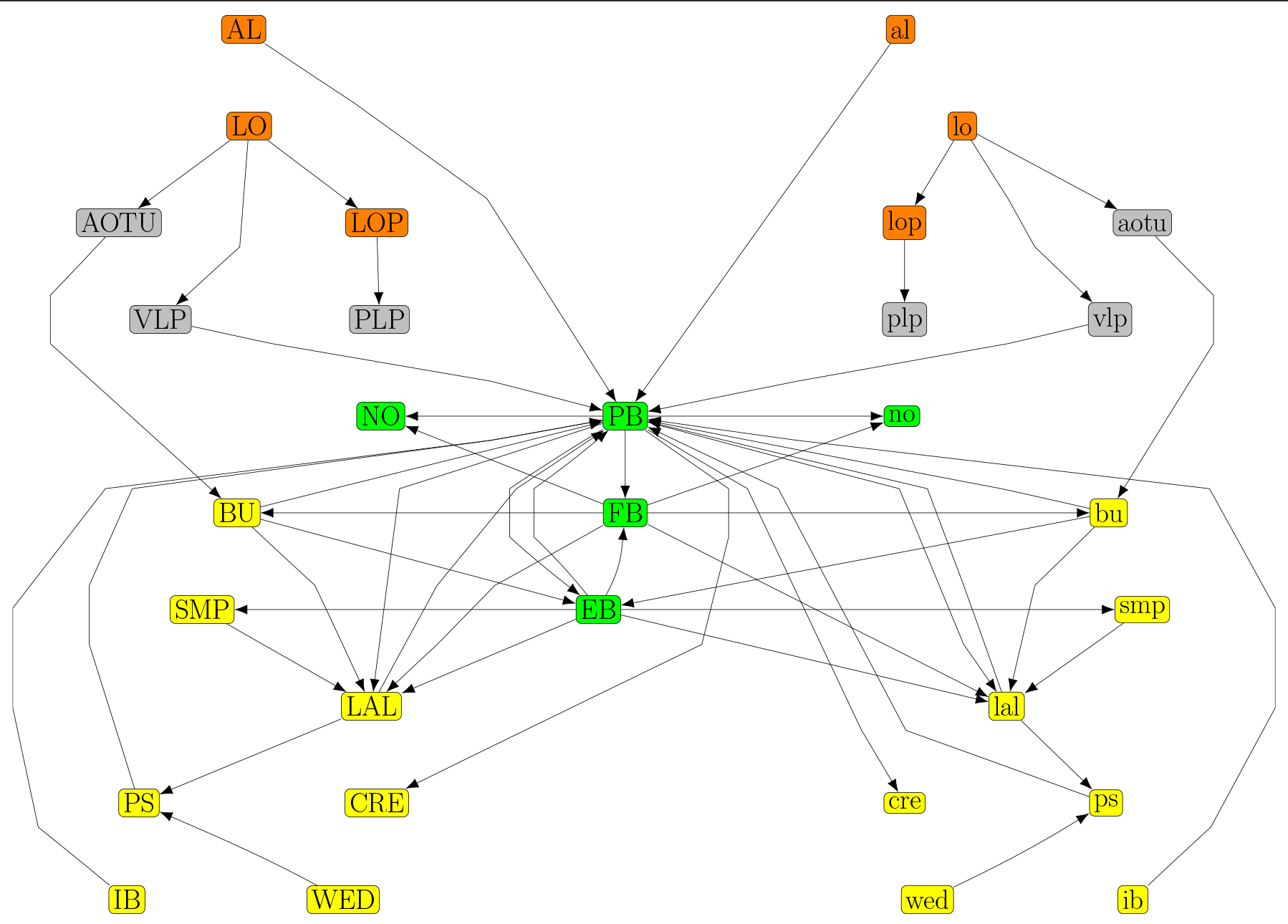
Information flow between CX neuropils (green), sensory neuropils (orange), neuropils that receive input from sensory neuropils (gray), and other accessory neuropils connected to the CX (yellow).

Spiking responses have been recorded from cells in FB during CX-related experiments using *Drosophila* [12, p. 64], from neurons supplying PB, tangential/pontine cells in FB, and ring cells in EB in *Neobellieria* [11], and from CX neurons in other insects [9].

## V. Identified Neurons in the Central Complex

In all neuropil innervation diagrams depicted below, arrow heads represent presynaptic arborizations and arrow tails represent postsynaptic arborizations.

### 5.1 Index of Identified Neurons

**Table II.** Identified local neurons in CX neuropils.

| Neuropil | References |
| --- | --- |
| FB | [15, p. 349, 352], [22, p. 1439] |
| PB | [1, p. 1743],[5, p. 1007] |

**Table III.** Identified projection neurons connecting CX and accessory neuropils. Additional inputs from vision neuropils and AOTU to ATL, BU, LAL, PLP, PS, SMP, VLP and outputs to locomotion neuropils have also been observed [25, p. 939], [24, Fig. 6], [1], [23, p. 9],.

| Neuron Family | Locations of Postsynaptic Arborizations (Dendrites) | Locations of Presynaptic Arborizations (Axons) | References |
| --- | --- | --- | --- |
| AL-PB | AL | PB | [15, p. 362, Fig. 24a] |
| BU-EB | BU | EB | [15, p. 352, Fig. 20a-d] |
| BU-LAL | BU | LAL | [15, Fig. 14] |
| DAL | SMP | LAL, SMP | [26, p. 680] |
| EB-FB | EB, FB | EB, FB | [15, Fig. 13a] |
| EB-FB-LAL-SMP | EB | FB, LAL, SMP | [15, p. 353, Fig. 21a-b] |
| EB-LAL-PB | EB | EB, LAL, PB | [1, Fig. 4J-M] |
| EB-NO | EB, NO | EB, NO | [15, Fig. 13b] |
| F | BU, FB, LAL, NO | FB, NO | [15, Fig. 14, 22, p. 353, 356] |
| FB-BU-LAL | FB | BU, LAL | [15, Fig. 14] |
| FB-NO | FB, NO | FB | [15, Fig. 11] |
| IB-LAL-PS-PB | IB, LAL, PS | PB | [1, p. 1743, Fig. 4A] [5, Fig. 3N] |
| PB-EB | PB | EB | [15, p. 350, Fig. 12b] |
| PB-EB-BU | PB | BU, EB | [15, Fig. 10b] |
| PB-EB-LAL | PB | EB, LAL | [1, Fig. 5E] |
| PB-EB-NO | PB | EB, NO | [1, p. 1745, Fig. 5G] |
| PB-FB | PB | FB, PB | [15, Fig. 12b] |
| PB-FB-CRE | PB | CRE, FB | [1, Fig. 6F] [5, Fig. 3L] |
| PB-FB-EB | EB, FB, PB | EB, FB | [15, Fig. 12e,f] |
| PB-FB-LAL | PB | FB, LAL | [1, Fig. 6F-H] |
| PB-FB-LAL-CRE | PB | CRE, FB, LAL | [5, Fig. 3M] |
| PB-FB-NO | PB | FB, NO | [1, p. 1746, Fig. 5L] |
| PS-IB-PB | IB, PS | PB | [5, Fig. 3S-T] |
| PS-PB | PS | PB | [5, Fig. 3R] |
| WED-PS-PB | PS, WED | PB | [1, p. 1744, Fig. 4B,D] |

**Table IV.** Projection neurons connecting CX and accessory neuropils with unresolved neurite types.

| Neuron Family | Locations of Arborizations | References |
| --- | --- | --- |
| CC | LAL, NO |  |
| EB-NO | EB, NO | [15, p. 351] |
| FB-EB | EB, FB | [15, p. 351] |
| FB-NO | FB, NO | [15, Fig. 11] |

### 5.1 Neurotransmitter Profiles

A range of neurotransmitters appear to be present in the CX neuropils (Table 5). Neurotransmitters associated with specific CX neural pathways have been identified (Table 6); however, the neurotransmitter associated with each specific arborization remains unclear.

**Table V.** Neurotransmitters in the fruit fly CX (adapted from [6]). Columnar neurons include those that connect PB to other neuropils or connect FB and EB, NO, LAL, or other neuropils. Tangential neurons include PB local neurons, F neurons in FB, and BU-EB neurons in EB. Ascending neurons connect the subesophageal ganglion to FB.

| Neurotransmitters | Neuropil | Cell Type | References |
| --- | --- | --- | --- |
| Acetylcholine | PB, FB, EB, NO | ? | [27] |
| Glutamate | PB, FB, EB, NO | Col, Tan | [28, 27] |
| GABA | FB, EB | Tan | [15] |
| Dopamine | PB, FB, EB, NO | Tan | [29] |
| Histamine | ? | ? | [30] |
| Octopamine | PB, FB | Asc, Tan | [31] |
| Serotonin | FB, EB, NO | Tan | [32] |
| Nitric oxide | FB, EB | ? | [33] |

**Table VI.** Neurotransmitter profiles of specific neural pathways in the fruit fly CX (adapted from [1, Fig. 7c] and [2]).

| Neuron Family | Neurotransmitter |
| --- | --- |
| AL-PB | ? |
| BU-EB | Acetylcholine [2, p.1598], Glutamate, GABA |
| BU-LAL | ? |
| DAL | ? |
| EB-FB | ? |
| EB-FB-LAL-SMP | Dopamine |
| EB-LAL-PB | Acetylcholine |
| EB-NO | ? |
| F | Dopamine |
| FB local | Acetylcholine |
| FB-BU-LAL | ? |
| FB-EB | ? |
| FB-NO | ? |
| IB-LAL-PS-PB | Dopamine |
| PB local | Acetylcholine, Glutamate |
| PB-EB | ? |
| PB-EB-BU | ? |
| PB-EB-LAL | Acetylcholine |
| PB-EB-NO | Acetylcholine |
| PB-FB | ? |
| PB-FB-CRE | ? |
| PB-FB-LAL | Acetylcholine |
| PB-FB-NO | Acetylcholine |
| PS-IB-PB | ? |
| PS-PB | ? |
| WED-PS-PB | Acetylcholine |

### 5.3 Local Neurons

#### 5.3.1 PB Local Neurons

Different studies of PB have identified 8 [5, p. 1007] or 10 [1, p. 1743] distinct local neurons. Table 7 and Fig. 12 assume the presence of 8 glomeruli on each side of PB as indicated by [5], that R2-R9 in [5] correspond to R1-R8 in [1], and that postsynaptic arborizations are spaced 7 glomeruli apart in all but the first 2 neuron types. The total number of each local neuron type is unclear.

**Table VII.** PB local neurons.

|  | Label |
| --- | --- |
| 1 | PB/L9/b-PB/L[4-8]/s |
| 2 | PB/L8\|R1\|R9/b-PB/L[1-9]\|R[1-9]/s |
| 3 | PB/L7\|R2/b-PB/L[1-9]\|R[1-9]/s |
| 4 | PB/L6\|R3/b-PB/L[1-9]\|R[1-9]/s |
| 5 | PB/L5\|R4/b-PB/L[1-9]\|R[1-9]/s |
| 6 | PB/R5\|L4/b-PB/R[1-9]\|L[1-9]/s |
| 7 | PB/R6\|L3/b-PB/R[1-9]\|L[1-9]/s |
| 8 | PB/R7\|L2/b-PB/R[1-9]\|L[1-9]/s |
| 9 | PB/R8\|L1\|L9/b-PB/R[1-9]\|L[1-9]/s |
| 10 | PB/R9/b-PB/R[4-8]/s |

**Fig. 12.**
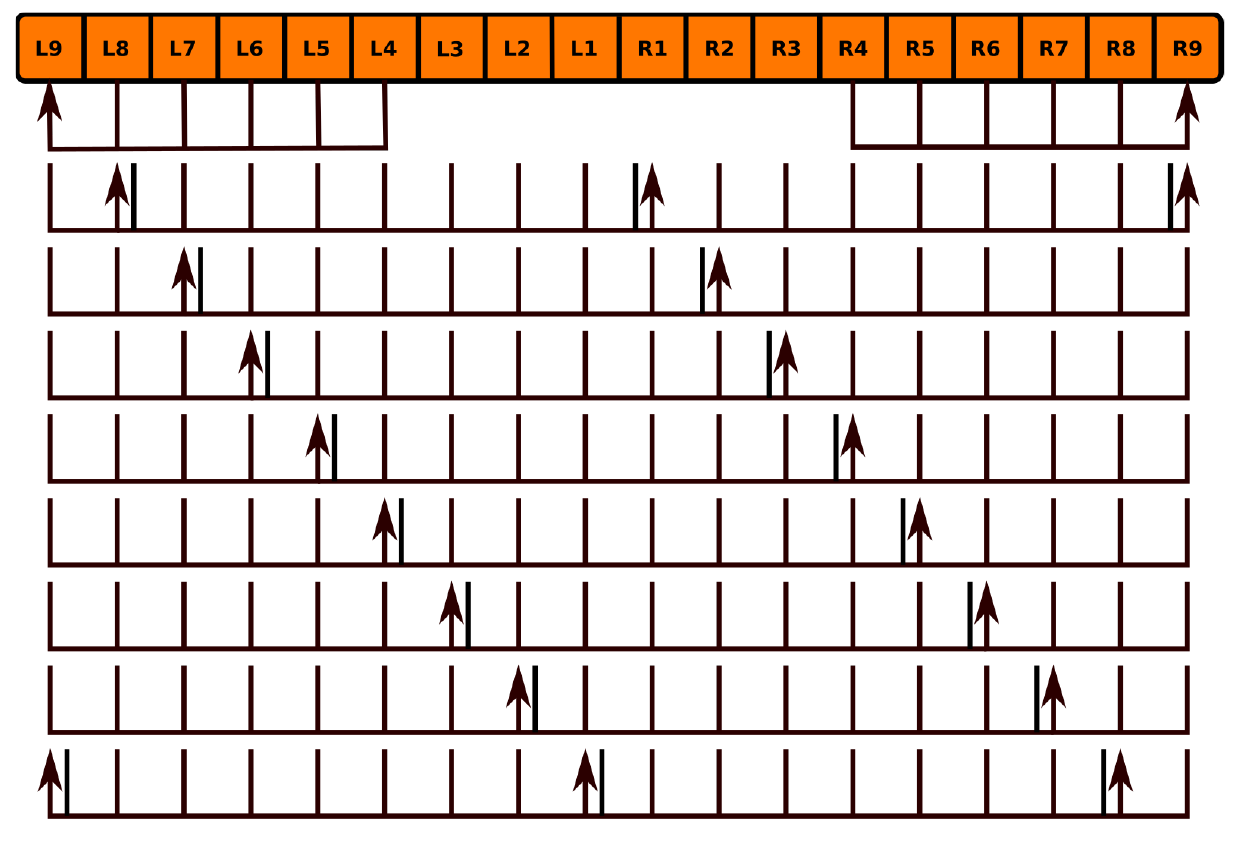
Innervation pattern of PB local neurons (Table 7).

### 5.3.2 FB Local Neurons

Several classes of local neurons referred to as pontine neurons have been observed to connect different regions of FB with each other [15, p. 349], [16, p. 1507]. One class (Table 8) comprises symmetric neurons that connect each segment on one side of the FB with each segment on the other side such that the presynaptic and postsynaptic arborizations are 4 segments apart [15, p. 352] (although more recent work suggests that each neuron might be a bundle of 2 neurons [22, p. 1439]). Judging by the structure of pontine neurons in other insects [34], arborizations might not be strictly confined to targeted regions. Other classes dorsoventrally connecting different layers in FB may exist, but they have not been systematically identified [15, p. 349]. It is unclear whether local neurons other than pontine neurons exist in the FB.

**Table VIII.** One identified class of FB local neurons.

|  | Label |
| --- | --- |
| 1 | FB/(3,L4)/s-FB/(3,R1)/b |
| 2 | FB/(3,L3)/s-FB/(3,R2)/b |
| 3 | FB/(3,L2)/s-FB/(3,R3)/b |
| 4 | FB/(3,L1)/s-FB/(3,R4)/b |
| 5 | FB/(3,R4)/s-FB/(3,L1)/b |
| 6 | FB/(3,R3)/s-FB/(3,L2)/b |
| 7 | FB/(3,R2)/s-FB/(3,L3)/b |
| 8 | FB/(3,R1)/s-FB/(3,L4)/b |

**Fig. 13.**
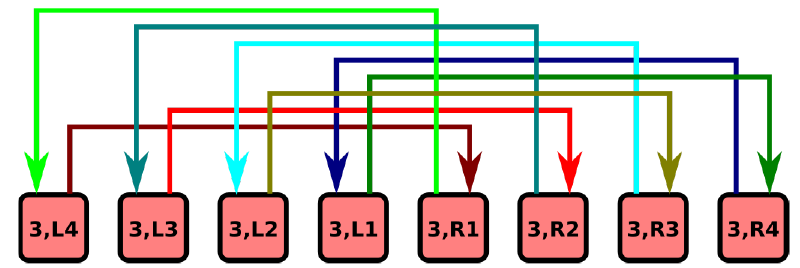
Innervation pattern of FB local neurons (Table 8).

### 5.3.3 EB Local Neurons

Although there appear to be local neurons in EB [3], they do not appear to have been systematically identified yet.

## 5.4 Projection Neurons

### 5.4.1 BU-EB Projection Neurons

Neurons with postsynaptic arborizations in BU and presynaptic arborizations in EB are typically referred to as ring or R neurons [15, p. 352] by virtue of the shape of their EB arborizations. 5 types of ring neurons (R1, R2, R3, R4m, R4d) have been observed [16, p. 1509]; specific ring neuron types appear to be essential to different visual behaviors [35, p. 120]. Each ring neuron type arborizes in a single microglomerulus [8, p. 262] and a different portion of the EB radius (Fig. 14); these types correspond to different sets of BU microglomeruli (and hence comprise multiple neurons). About 20 of each of these types of neurons have been estimated in each hemisphere of the fruit fly brain [16, p. 1510]; combined with visual confirmation of the presence of 80 microglomeruli (§ 3.5), this suggests that there are 16 of each neuron type present in BU. Some ring neurons are GABAergic [1, p. 1750], while others are glutamatergic [1, Fig. 7C]; their synaptic connections to other neurons in EB therefore seem to be inhibitory. There is recent evidence that some ring neurons may be cholinergic and hence possess excitatory synapses [2, p. 1598]. Coincident synapses (i.e., those in which two independent presynaptic zones coincide with a single postsynaptic zone) have been observed in EB between ring neurons in specific domains and other neurons both in and outside of those domains. [2, p. 1592]; it seems that such synapses may also exist between other neurons that innervate EB [2, p. 1594]. Connections between AOTU and BU have been observed [23, p. 9]; these presumably constitute a pathway for input visual information from LO via AOTU [25, p. 939] to EB via ring neurons.

**Fig. 14.**
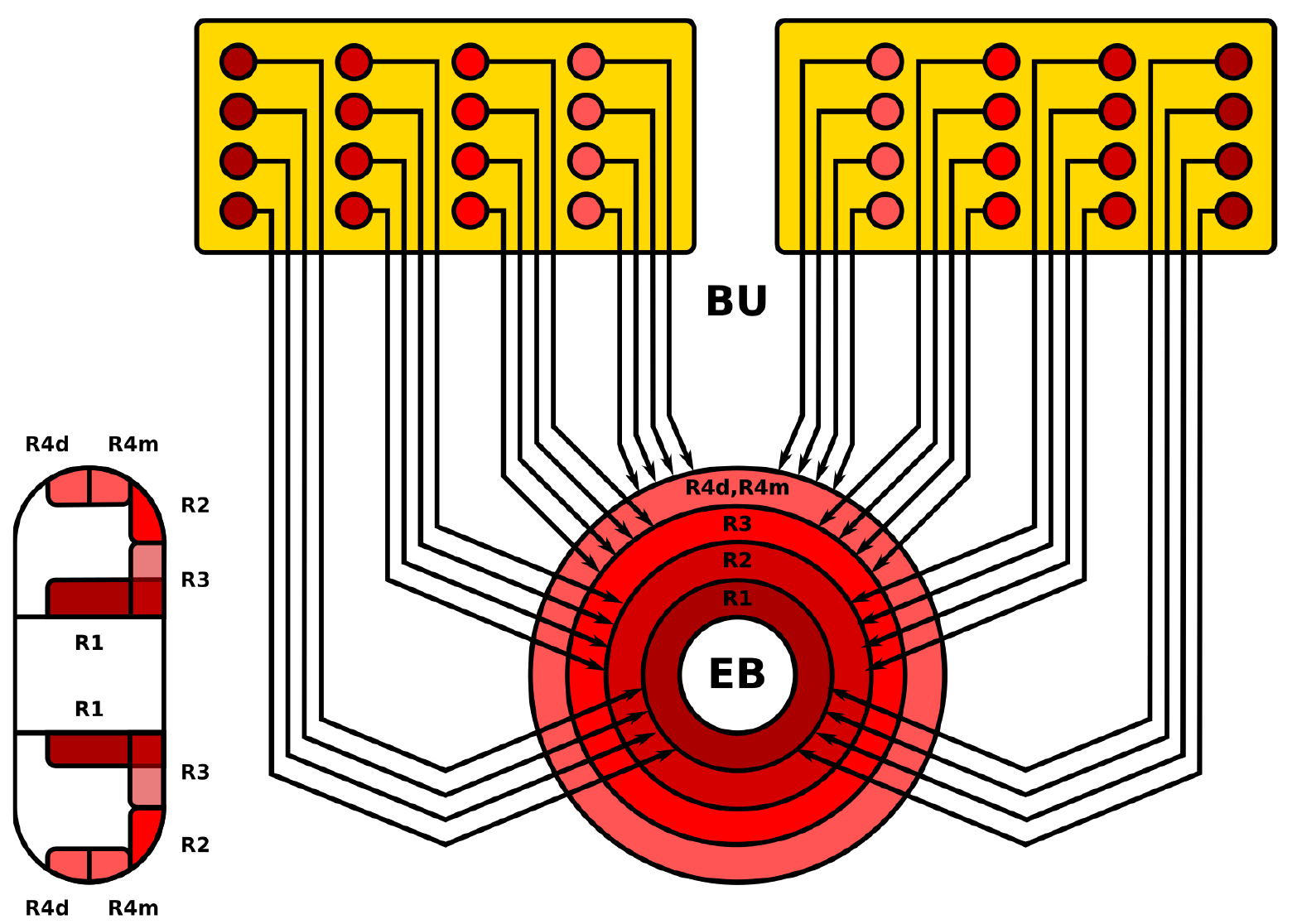
Spatial organization of ring neuron arborizations in BU microglomeruli (yellow) and regions in EB (red); posterior is left of sagittal section of EB, anterior is right. The depicted EB regions do not exactly correspond to the rings in Fig. 6c. Only a fraction of the microglomeruli/neurons are depicted. The spatial organization of each neuron type’s microglomeruli is unclear.

### 5.4.2 EB-FB-LAL-SMP Projection Neurons

Neurons with presynaptic ring-shaped arborizations in EB and postsynaptic arborizations in other neuropils are referred to as extrinsic ring neurons. Two types (ExR1, ExR2) have been observed; these neurons appear to constitute a dopaminergic pathway [1, Fig. 7C]. ExR1 neurons are presynaptic in EB, FB, and LAL, and postsynaptic in SMP [15, p. 353] [VirtualFlyBrain]. ExR2 appears to have presynaptic arborizations in EB and arborizations in SMP, but the remainder of the neuron’s structure has not been reconstructed.

### 5.4.3 EB-LAL-PB Projection Neurons

These neurons correspond to the EB-PB-VBO or EIP neurons in [1, Fig. 2]; they constitute a cholinergic pathway [1, Fig. 7C]. The neuron arborizations in Table 9 and Fig. 15 make the assumption that the C, O, and P rings in [1] collectively correspond to the P and M shells in [5] and that the C and P rings collectively correspond to the P shell.

**Table IX.** EB-LAL-PB neurons.

|  | Label |
| --- | --- |
| 1 | EB/(L8,P,[1-4])/s-EB/(L8,P,[1-4])/b-lal/RDG/b-PB/L9/b |
| 2 | EB/([L7,R8],[P,M],[1-4])/s-EB/(L8,[P,M],[1-4])/b-LAL/LDG/b-PB/R1\|L1/b |
| 3 | EB/([L8,L6],[P,M],[1-4])/s-EB/(L7,[P,M],[1-4])/b-lal/RVG/b-PB/L8/b |
| 4 | EB/([L5,L7],[P,M],[1-4])/s-EB/(L6,[P,M],[1-4])/b-LAL/LVG/b-PB/R2/b |
| 5 | EB/([L6,L4],[P,M],[1-4])/s-EB/(L5,[P,M],[1-4])/b-lal/RDG/b-PB/L7/b |
| 6 | EB/([L3,L5],[P,M],[1-4])/s-EB/(L4,[P,M],[1-4])/b-LAL/LDG/b-PB/R3/b |
| 7 | EB/([L4,L2],[P,M],[1-4])/s-EB/(L3,[P,M],[1-4])/b-lal/RVG/b-PB/L6/b |
| 8 | EB/([L1,L3],[P,M],[1-4])/s-EB/(L2,[P,M],[1-4])/b-LAL/LVG/b-PB/R4/b |
| 9 | EB/([L2,R1],[P,M],[1-4])/s-EB/(L1,[P,M],[1-4])/b-lal/RDG/b-PB/L5/b |
| 10 | EB/([R2,L1],[P,M],[1-4])/s-EB/(R1,[P,M],[1-4])/b-LAL/LDG/b-PB/R5/b |
| 11 | EB/([R1,R3],[P,M],[1-4])/s-EB/(R2,[P,M],[1-4])/b-lal/RVG/b-PB/L4/b |
| 12 | EB/([R4,R2],[P,M],[1-4])/s-EB/(R3,[P,M],[1-4])/b-LAL/LVG/b-PB/R6/b |
| 13 | EB/([R3,R5],[P,M],[1-4])/s-EB/(R4,[P,M],[1-4])/b-lal/RDG/b-PB/L3/b |
| 14 | EB/([R6,R4],[P,M],[1-4])/s-EB/(R5,[P,M],[1-4])/b-LAL/LDG/b-PB/R7/b |
| 15 | EB/([R5,R7],[P,M],[1-4])/s-EB/(R6,[P,M],[1-4])/b-lal/RVG/b-PB/L2/b |
| 16 | EB/([R8,R6],[P,M],[1-4])/s-EB/(R7,[P,M],[1-4])/b-LAL/LVG/b-PB/R8/b |
| 17 | EB/([R7,L8],[P,M],[1-4])/s-EB/(R8,[P,M],[1-4])/b-lal/RDG/b-PB/L1\|R1/b |
| 18 | EB/(R8,P,[1-4])/s-EB/(R8,P,[1-4])/b-LAL/LDG/b-PB/R9/b |

**Fig. 15.**
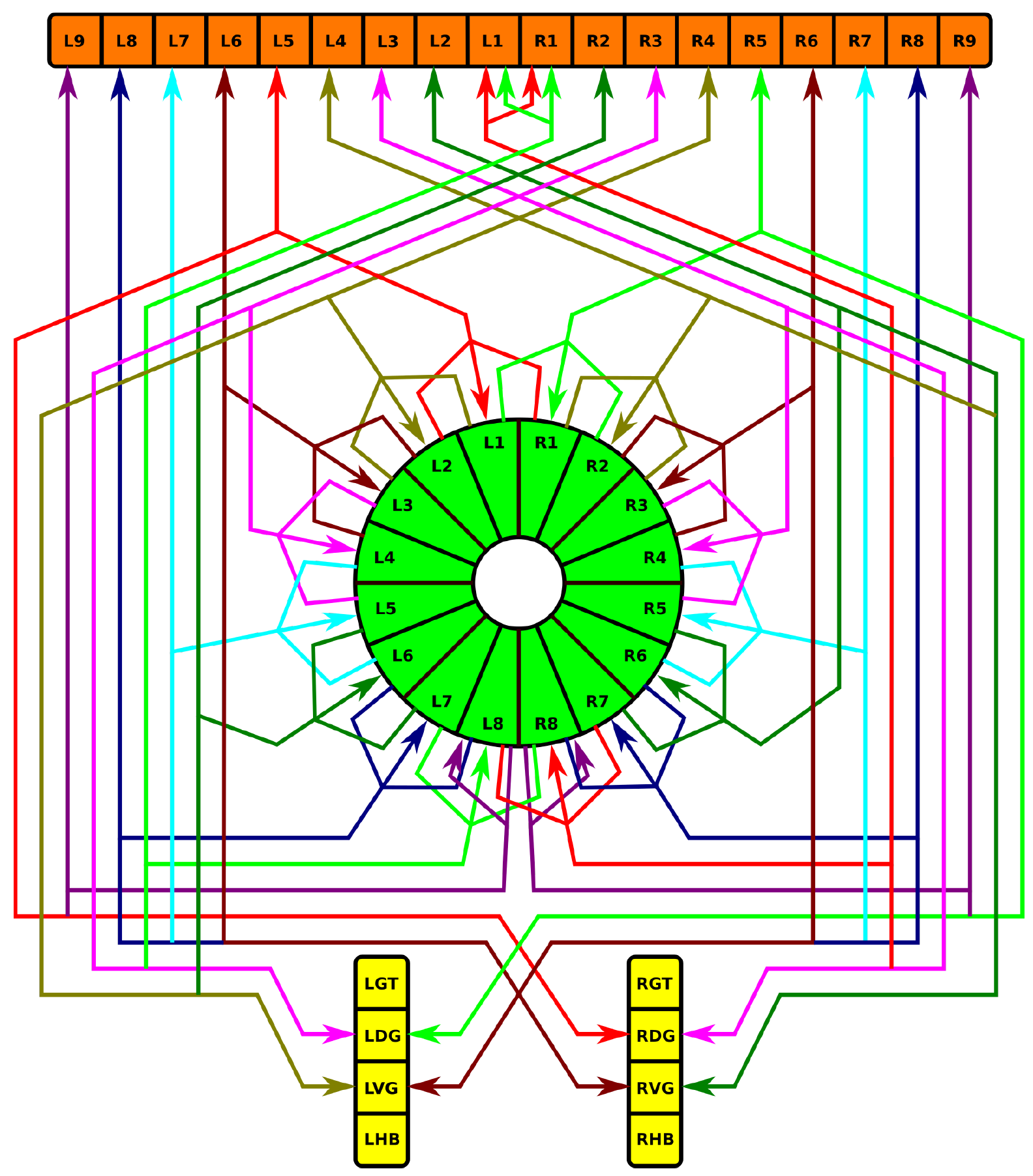
Neuropil innervation pattern for EB-LAL-PB neurons (Table 9).

### 5.4.4 F Projection Neurons

F neurons innervate entire layers of FB, with some types associated with specific layers [15, p. 353]. Fm neurons have bleb-like (presynaptic) arborizations in layer 2 of FB; Fm1 neurons have spiny (postsynaptic arborizations in SLP or SIP, Fm2 neurons have spiny arborizations in LAL, and Fm3 neurons have spiny arborizations in ICL [15, p. 353]. Some F1 neurons have spiny branches in LAL [15, p. 354], while other F1 neurons have spiny branches that innervate the entire BU [15, p. 354].

### 5.4.5 IB-LAL-PS-PB Projection Neurons

These neurons corresponds to the CVLP-IDFP-VMP-PB or CIVP neurons in [1, Fig. 2] and the PB-LAL-PS neurons in [5, Fig. 3N]. They receive indirect input from the vision system and innervate all glomeruli in PB indicated in Table 10:

**Table X.** IB-LAL-PS-PB neurons.

|  | Label |
| --- | --- |
| 1 | IB/L/s-LAL/LHB/s-PS/L/s-PB/L[2-9]\|R[2-9]/b |
| 2 | ib/R/s-lal/RHB/s-ps/R/s-PB/L[2-9]\|R[2-9]/b |

**Fig. 16.**
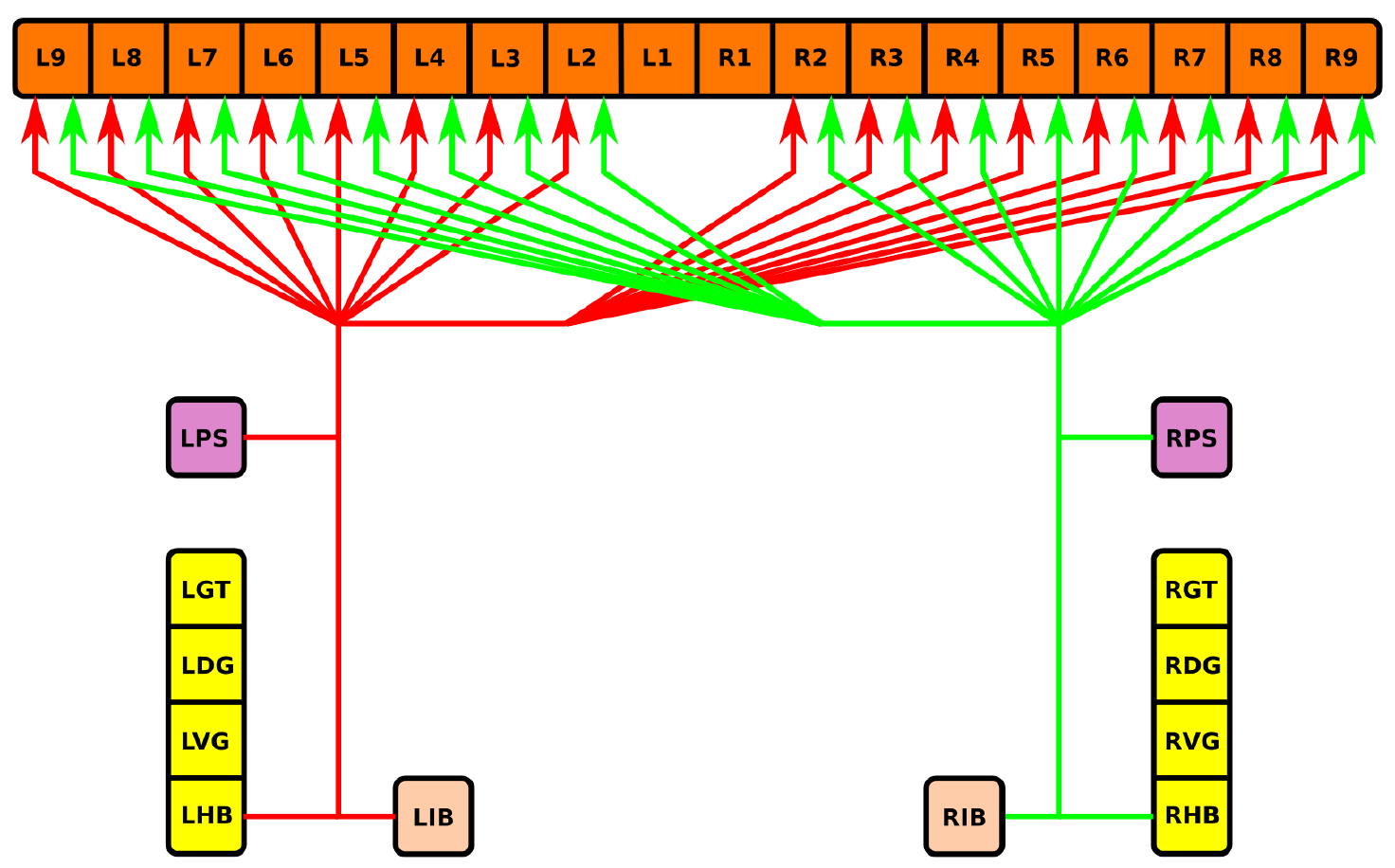
Neuropil innervation pattern for IB-LAL-PS-PB neurons (Table 10).

### 5.4.6 PB-EB-BU Projection Neurons

Neurons with postsynaptic arborizations in PB and presynaptic arborizations in EB and BU have been observed. These correspond to the PB-EB-LTR neurons in [15, Fig. 10b]. Details regarding possible neuron types have not been determined.

### 5.4.7 PB-EB-NO Projection Neurons

Neurons with postsynaptic arborizations in PB and presynaptic arborizations in EB and NO are referred to as PEN neurons in [1, p. 1745]. The constitute a cholinergic pathway [1, Fig. 7C]. The connections in Table 11 and Fig. 17 are based upon [5, Fig. 14a], which differ from those described in [1, Fig. 5g].

**Table XI.** PB-EB-NO neurons.

|  | Label |
| --- | --- |
| 1 | PB/L9/s-EB/6/b-no/(1,R)/b |
| 2 | PB/L8/s-EB/7/b-no/(1,R)/b |
| 3 | PB/L7/s-EB/8/b-no/(1,R)/b |
| 4 | PB/L6/s-EB/1/b-no/(1,R)/b |
| 5 | PB/L5/s-EB/2/b-no/(1,R)/b |
| 6 | PB/L4/s-EB/3/b-no/(1,R)/b |
| 7 | PB/L3/s-EB/4/b-no/(1,R)/b |
| 8 | PB/L2/s-EB/5/b-no/(1,R)/b |
| 9 | PB/R2/s-EB/5/b-NO/(1,L)/b |
| 10 | PB/R3/s-EB/6/b-NO/(1,L)/b |
| 11 | PB/R4/s-EB/7/b-NO/(1,L)/b |
| 12 | PB/R5/s-EB/8/b-NO/(1,L)/b |
| 13 | PB/R6/s-EB/1/b-NO/(1,L)/b |
| 14 | PB/R7/s-EB/2/b-NO/(1,L)/b |
| 15 | PB/R8/s-EB/3/b-NO/(1,L)/b |
| 16 | PB/R9/s-EB/4/b-NO/(1,L)/b |

**Fig. 17.**
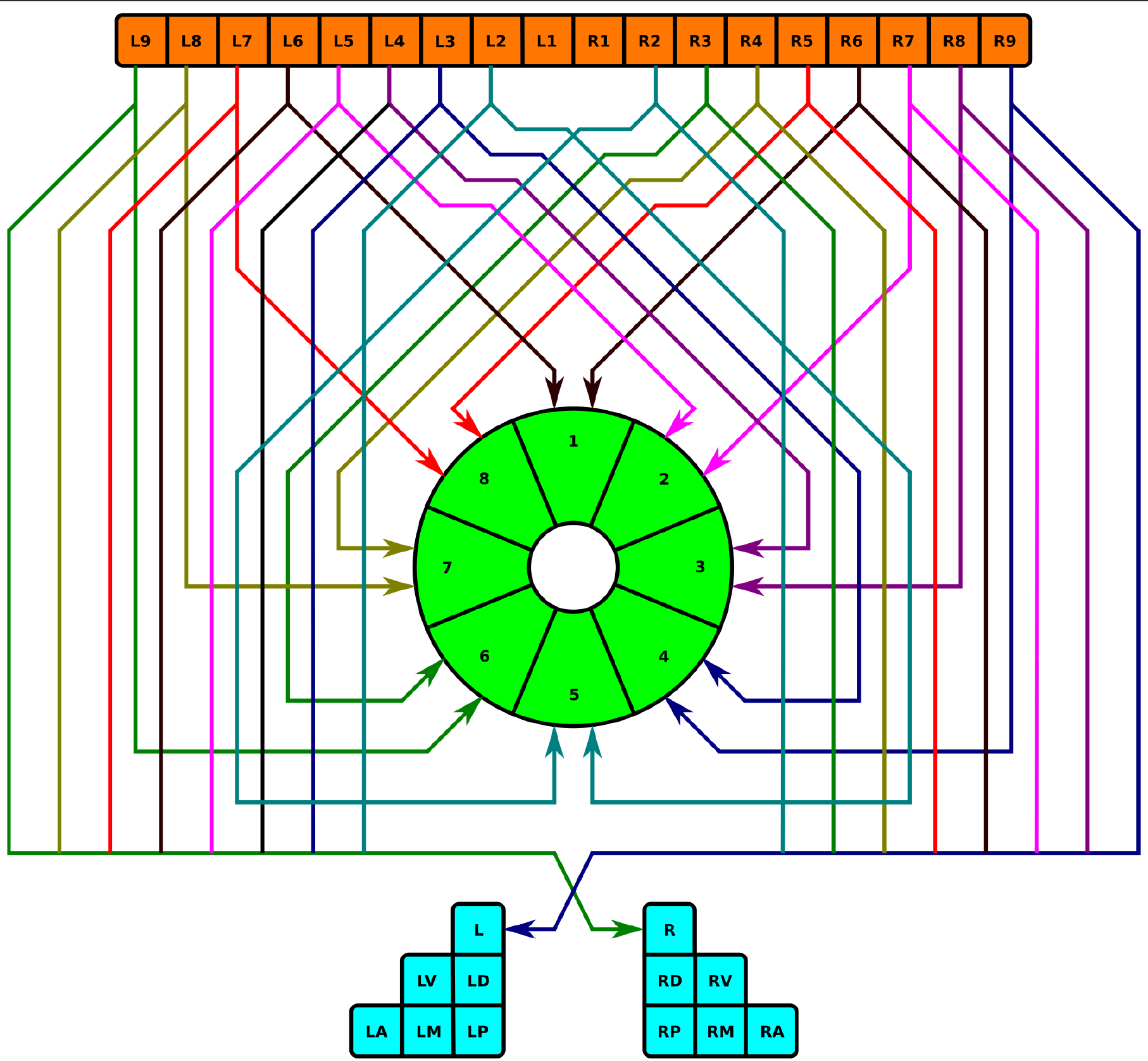
Neuropil innervation pattern for PB-EB-NO neurons (Table 11).

### 5.4.8 PB-EB-LAL Projection Neurons

These neurons have postsynaptic arborizations in PB and presynaptic arborizations in EB and LAL; they correspond to PB-EB-IDFP or PEI neurons in [1, Fig. 2]. These neurons constitute a cholinergic pathway [1, Fig. 7C]. The connections in Table 12 and Fig. 18 are based upon [5, Fig. 14b], which differ from those described in [1, Fig. 5e].

**Table XII.** PB-EB-LAL neurons.

|  | Label |
| --- | --- |
| 1 | PB/L8/s-EB/6/b-lal/RVG/b |
| 2 | PB/L7/s-EB/7/b-lal/RDG/b |
| 3 | PB/L6/s-EB/8/b-lal/RVG/b |
| 4 | PB/L5/s-EB/1/b-lal/RDG/b |
| 5 | PB/L4/s-EB/2/b-lal/RVG/b |
| 6 | PB/L3/s-EB/3/b-lal/RDG/b |
| 7 | PB/L2/s-EB/4/b-lal/RVG/b |
| 8 | PB/L1/s-EB/5/b-lal/RDG/b |
| 9 | PB/R1/s-EB/5/b-LAL/LDG/b |
| 10 | PB/R2/s-EB/4/b-LAL/LVG/b |
| 11 | PB/R3/s-EB/3/b-LAL/LDG/b |
| 12 | PB/R4/s-EB/2/b-LAL/LVG/b |
| 13 | PB/R5/s-EB/1/b-LAL/LDG/b |
| 14 | PB/R6/s-EB/8/b-LAL/LVG/b |
| 15 | PB/R7/s-EB/7/b-LAL/LDG/b |
| 16 | PB/R8/s-EB/6/b-LAL/LVG/b |

**Fig. 18.**
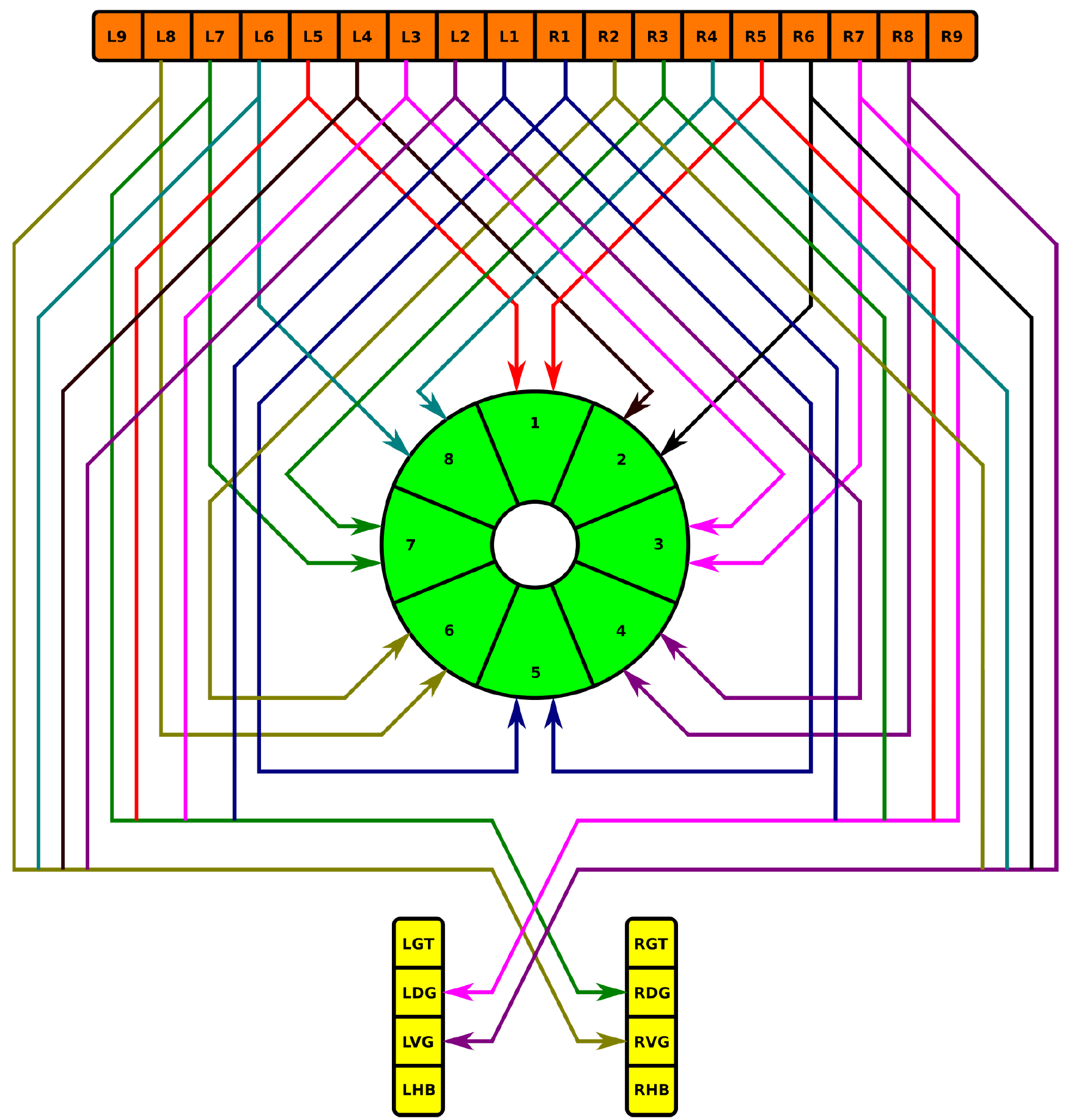
Neuropil innervation pattern for PB-EB-LAL neurons (Table 12).

### 5.4.9 PB-FB-CRE Projection Neurons

The neurons in Table 13 and Fig. 19 correspond to the PB-FB-VBO or EIP neurons that connect to RB in [1, Fig. 2]. They constitute a cholinergic pathway [1, Fig. 7C].

**Table XIII.** PB-FB-CRE neurons.

|  | Label |
| --- | --- |
| 1 | PB/L1/s-FB/([3-4],L4)/s-CRE/LRB/b |
| 2 | PB/R1/s-FB/([3-4],L4)/s-CRE/LRB/b |
| 3 | PB/R2/s-FB/([3-4],L3)/s-CRE/LRB/b |
| 4 | PB/R3/s-FB/([3-4],L2)/s-CRE/LRB/b |
| 5 | PB/R4/s-FB/([3-4],L1)/s-CRE/LRB/b |
| 6 | PB/R4/s-FB/([3-4],R1)/s-CRE/LRB/b |
| 7 | PB/R5/s-FB/([3-4],R2)/s-CRE/LRB/b |
| 8 | PB/R6/s-FB/([3-4],R3)/s-CRE/LRB/b |
| 9 | PB/R7/s-FB/([3-4],R4)/s-CRE/LRB/b |
| 10 | PB/L7/s-FB/([3-4],L4)/s-cre/RRB/b |
| 11 | PB/L6/s-FB/([3-4],L3)/s-cre/RRB/b |
| 12 | PB/L5/s-FB/([3-4],L2)/s-cre/RRB/b |
| 13 | PB/L4/s-FB/([3-4],L1)/s-cre/RRB/b |
| 14 | PB/L4/s-FB/([3-4],R1)/s-cre/RRB/b |
| 15 | PB/L3/s-FB/([3-4],R2)/s-cre/RRB/b |
| 16 | PB/L2/s-FB/([3-4],R3)/s-cre/RRB/b |
| 17 | PB/L1/s-FB/([3-4],R4)/s-cre/RRB/b |
| 18 | PB/R1/s-FB/([3-4],R4)/s-cre/RRB/b |

**Fig. 19.**
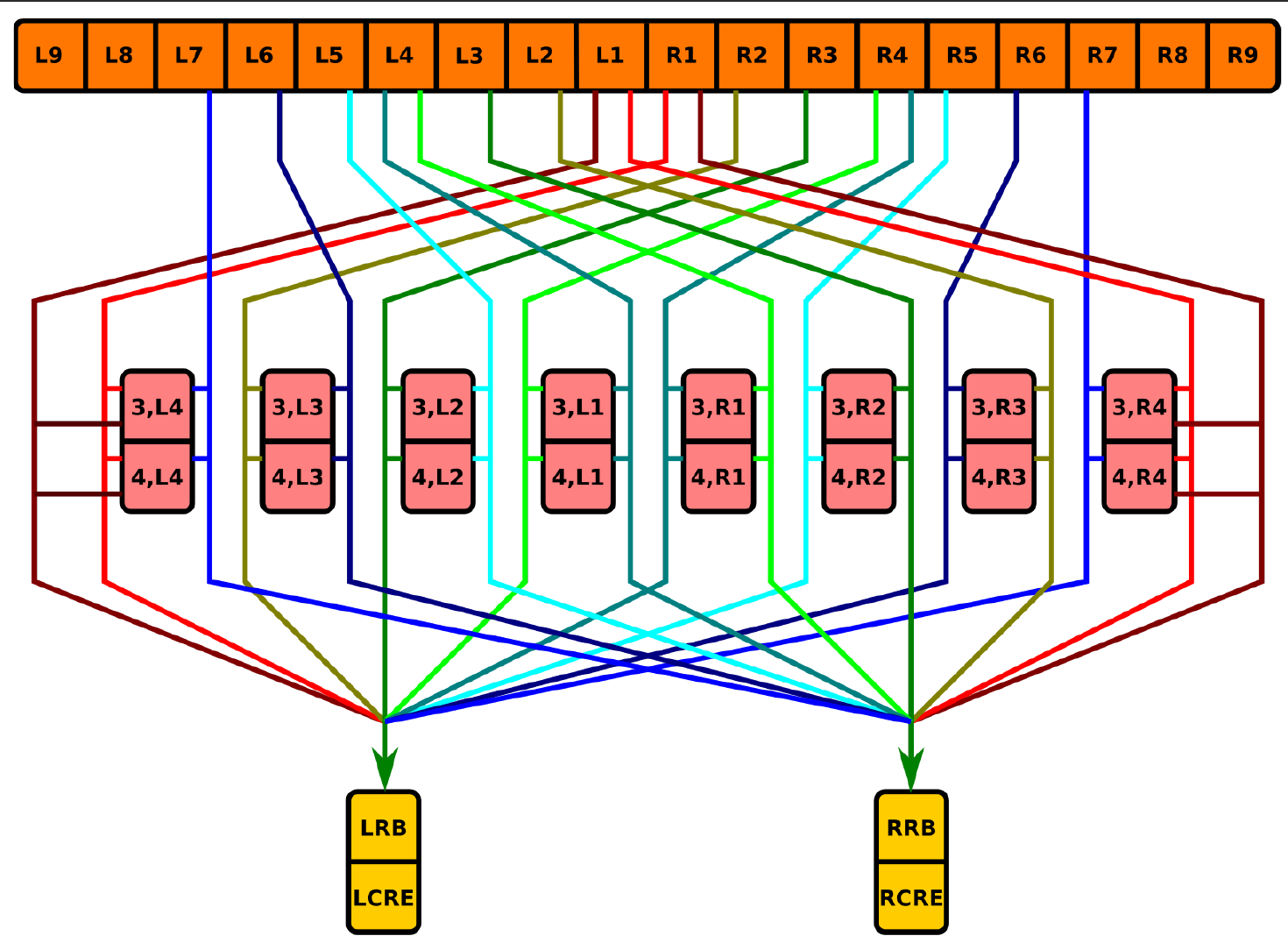
Neuropil innervation pattern for PB-FB-CRE neurons (Table 13).

### 5.4.10 PB-FB-NO Projection Neurons

Neurons with postsynaptic arborizations in PB and presynaptic arborizations in FB and NO are referred to as the vertical fiber system [15, Fig. 5b]; they correspond to the PFN neurons described in [1, p. 1745]. These neurons constitute a cholinergic pathway [1, Fig. 7C]. The 5 sets of PB-FB-NO neurons are listed in Tabs. 14, 15, 16, 17, and 18 and depicted in Fig. 20.

**Table XIV.** PB-FB-NO neurons innervating region (3,P) of NO.

|  | Label |
| --- | --- |
| 1 | PB/L2/s-FB/(1,R4)/b-no/(3,RP)/b |
| 2 | PB/L3/s-FB/(1,R4)/b-no/(3,RP)/b |
| 3 | PB/L4/s-FB/(1,R3)/b-no/(3,RP)/b |
| 4 | PB/L5/s-FB/(1,R2)/b-no/(3,RP)/b |
| 5 | PB/L6/s-FB/(1,R1)\|(1,L1)/b-no/(3,RP)/b |
| 6 | PB/L7/s-FB/(1,L2)/b-no/(3,RP)/b |
| 7 | PB/L8/s-FB/(1,L3)/b-no/(3,RP)/b |
| 8 | PB/L9/s-FB/(1,L4)/b-no/(3,RP)/b |
| 9 | PB/R2/s-FB/(1,L4)/b-NO/(3,LP)/b |
| 10 | PB/R3/s-FB/(1,L4)/b-NO/(3,LP)/b |
| 11 | PB/R4/s-FB/(1,L3)/b-NO/(3,LP)/b |
| 12 | PB/R5/s-FB/(1,L2)/b-NO/(3,LP)/b |
| 13 | PB/R6/s-FB/(1,L1)\|(1,R1)/b-NO/(3,LP)/b |
| 14 | PB/R7/s-FB/(1,R2)/b-NO/(3,LP)/b |
| 15 | PB/R8/s-FB/(1,R3)/b-NO/(3,LP)/b |
| 16 | PB/R9/s-FB/(1,R4)/b-NO/(3,LP)/b |

**Table XV.**
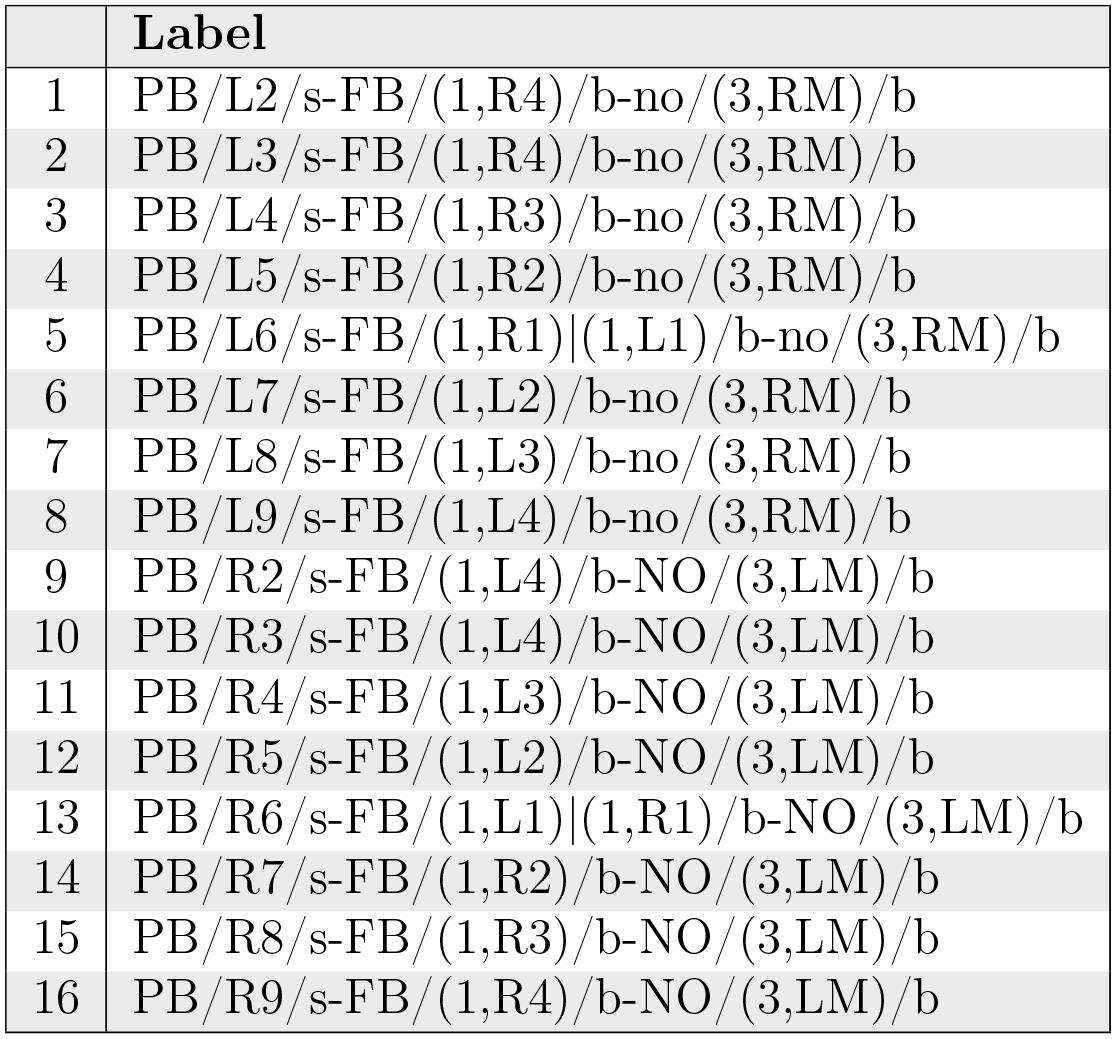
PB-FB-NO neurons innervating region (3,M) of NO

|  | Label |
| --- | --- |
| 1 | PB/L2/s-FB/(1,R4)/b-no/(3,RM)/b |
| 2 | PB/L3/s-FB/(1,R4)/b-no/(3,RM)/b |
| 3 | PB/L4/s-FB/(1,R3)/b-no/(3,RM)/b |
| 4 | PB/L5/s-FB/(1,R2)/b-no/(3,RM)/b |
| 5 | PB/L6/s-FB/(1,R1)\|(1,L1)/b-no/(3,RM)/b |
| 6 | PB/L7/s-FB/(1,L2)/b-no/(3,RM)/b |
| 7 | PB/L8/s-FB/(1,L3)/b-no/(3,RM)/b |
| 8 | PB/L9/s-FB/(1,L4)/b-no/(3,RM)/b |
| 9 | PB/R2/s-FB/(1,L4)/b-NO/(3,LM)/b |
| 10 | PB/R3/s-FB/(1,L4)/b-NO/(3,LM)/b |
| 11 | PB/R4/s-FB/(1,L3)/b-NO/(3,LM)/b |
| 12 | PB/R5/s-FB/(1,L2)/b-NO/(3,LM)/b |
| 13 | PB/R6/s-FB/(1,L1)\|(1,R1)/b-NO/(3,LM)/b |
| 14 | PB/R7/s-FB/(1,R2)/b-NO/(3,LM)/b |
| 15 | PB/R8/s-FB/(1,R3)/b-NO/(3,LM)/b |
| 16 | PB/R9/s-FB/(1,R4)/b-NO/(3,LM)/b |

**Table XVI.** PB-FB-NO neurons innervating region (3,A) of NO.

|  | Label |
| --- | --- |
| 1 | PB/L2/s-FB/(2,R4)/b-no/(3,RA)/b |
| 2 | PB/L3/s-FB/(2,R4)/b-no/(3,RA)/b |
| 3 | PB/L4/s-FB/(2,R3)/b-no/(3,RA)/b |
| 4 | PB/L5/s-FB/(2,R2)/b-no/(3,RA)/b |
| 5 | PB/L6/s-FB/(2,R1)\|(1,L1)/b-no/(3,RA)/b |
| 6 | PB/L7/s-FB/(2,L2)/b-no/(3,RA)/b |
| 7 | PB/L8/s-FB/(2,L3)/b-no/(3,RA)/b |
| 8 | PB/L9/s-FB/(2,L4)/b-no/(3,RA)/b |
| 9 | PB/R2/s-FB/(2,L4)/b-NO/(3,LA)/b |
| 10 | PB/R3/s-FB/(2,L4)/b-NO/(3,LA)/b |
| 11 | PB/R4/s-FB/(2,L3)/b-NO/(3,LA)/b |
| 12 | PB/R5/s-FB/(2,L2)/b-NO/(3,LA)/b |
| 13 | PB/R6/s-FB/(2,L1)\|(1,R1)/b-NO/(3,LA)/b |
| 14 | PB/R7/s-FB/(2,R2)/b-NO/(3,LA)/b |
| 15 | PB/R8/s-FB/(2,R3)/b-NO/(3,LA)/b |
| 16 | PB/R9/s-FB/(2,R4)/b-NO/(3,LA)/b |

**Table XVII.** PB-FB-NO neurons innervating region (2,D) of NO.

|  | Label |
| --- | --- |
| 1 | PB/L2/s-FB/(3,R4)/b-no/(2,RD)/b |
| 2 | PB/L3/s-FB/(3,R4)/b-no/(2,RD)/b |
| 3 | PB/L4/s-FB/(3,R3)/b-no/(2,RD)/b |
| 4 | PB/L5/s-FB/(3,R2)/b-no/(2,RD)/b |
| 5 | PB/L6/s-FB/(3,R1)\|(1,L1)/b-no/(2,RD)/b |
| 6 | PB/L7/s-FB/(3,L2)/b-no/(2,RD)/b |
| 7 | PB/L8/s-FB/(3,L3)/b-no/(2,RD)/b |
| 8 | PB/L9/s-FB/(3,L4)/b-no/(2,RD)/b |
| 9 | PB/R2/s-FB/(3,L4)/b-NO/(2,LD)/b |
| 10 | PB/R3/s-FB/(3,L4)/b-NO/(2,LD)/b |
| 11 | PB/R4/s-FB/(3,L3)/b-NO/(2,LD)/b |
| 12 | PB/R5/s-FB/(3,L2)/b-NO/(2,LD)/b |
| 13 | PB/R6/s-FB/(3,L1)\|(1,R1)/b-NO/(2,LD)/b |
| 14 | PB/R7/s-FB/(3,R2)/b-NO/(2,LD)/b |
| 15 | PB/R8/s-FB/(3,R3)/b-NO/(2,LD)/b |
| 16 | PB/R9/s-FB/(3,R4)/b-NO/(2,LD)/b |

**Table XVIII.**
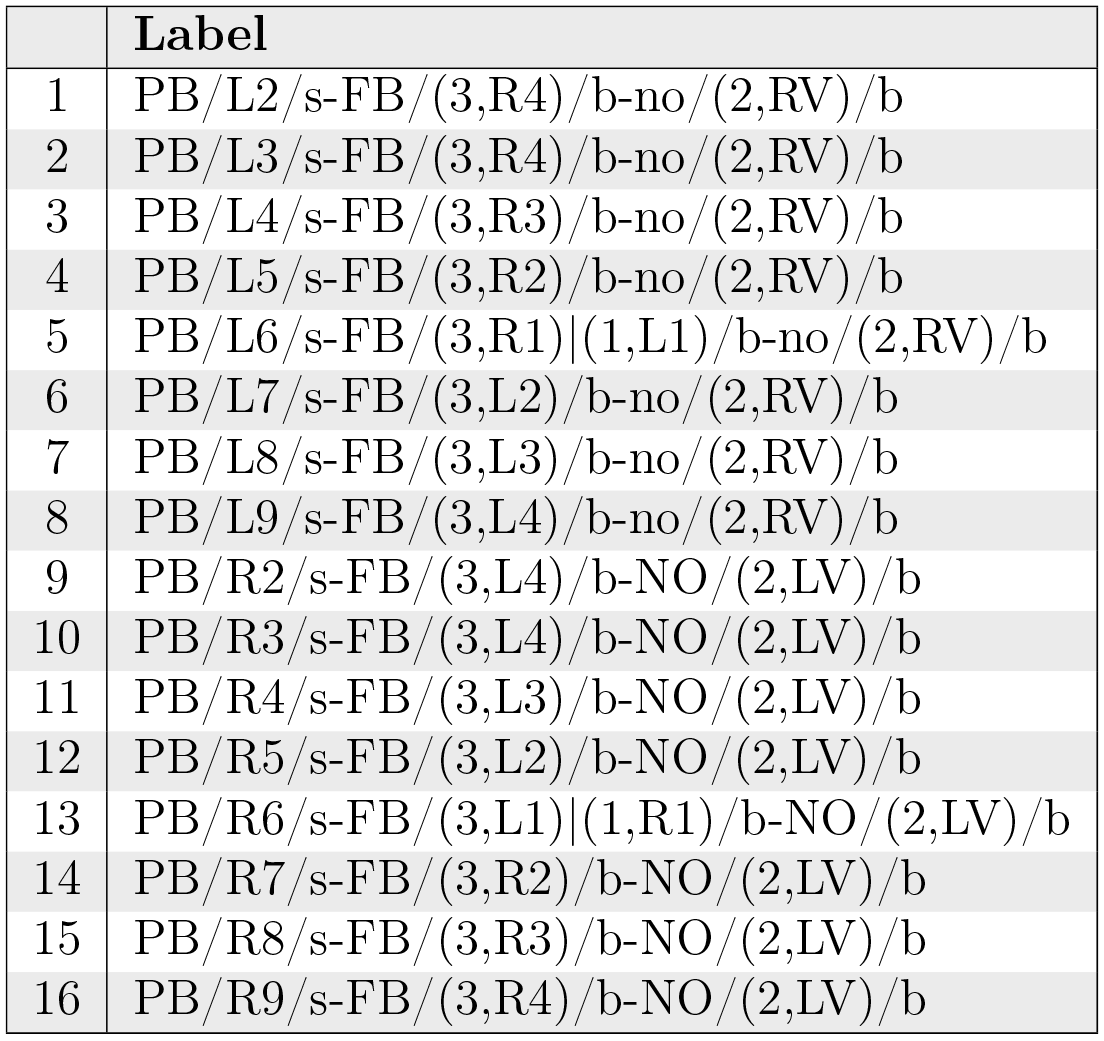
PB-FB-NO neurons innervating region (2,V) of NO.

|  | Label |
| --- | --- |
| 1 | PB/L2/s-FB/(3,R4)/b-no/(2,RV)/b |
| 2 | PB/L3/s-FB/(3,R4)/b-no/(2,RV)/b |
| 3 | PB/L4/s-FB/(3,R3)/b-no/(2,RV)/b |
| 4 | PB/L5/s-FB/(3,R2)/b-no/(2,RV)/b |
| 5 | PB/L6/s-FB/(3,R1)\|(1,L1)/b-no/(2,RV)/b |
| 6 | PB/L7/s-FB/(3,L2)/b-no/(2,RV)/b |
| 7 | PB/L8/s-FB/(3,L3)/b-no/(2,RV)/b |
| 8 | PB/L9/s-FB/(3,L4)/b-no/(2,RV)/b |
| 9 | PB/R2/s-FB/(3,L4)/b-NO/(2,LV)/b |
| 10 | PB/R3/s-FB/(3,L4)/b-NO/(2,LV)/b |
| 11 | PB/R4/s-FB/(3,L3)/b-NO/(2,LV)/b |
| 12 | PB/R5/s-FB/(3,L2)/b-NO/(2,LV)/b |
| 13 | PB/R6/s-FB/(3,L1)\|(1,R1)/b-NO/(2,LV)/b |
| 14 | PB/R7/s-FB/(3,R2)/b-NO/(2,LV)/b |
| 15 | PB/R8/s-FB/(3,R3)/b-NO/(2,LV)/b |
| 16 | PB/R9/s-FB/(3,R4)/b-NO/(2,LV)/b |

**Fig. 20.**
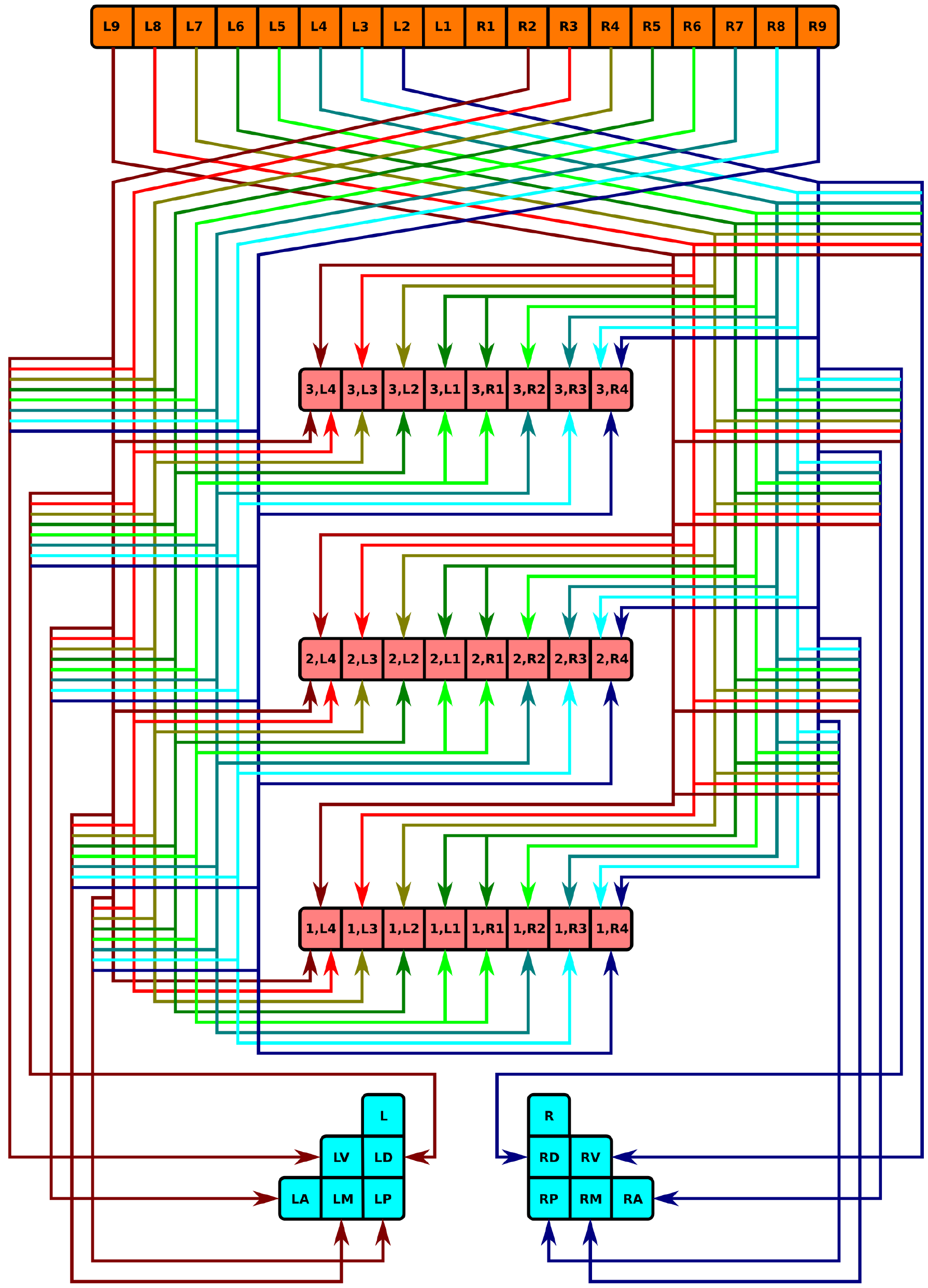
Neuropil innervation pattern for PB-FB-NO neurons (Tabs. 14, 15, 16, 17, 18).

### 5.4.11 PB-FB-LAL Projection Neurons

Neurons with postsynaptic arborizations in PB and FB and presynaptic arborizations in LAL are referred to as EIP neurons in [1, p. 1745]. They correspond to the PB-FB-VBO or EIP neurons that connect to HB in [1, Fig. 2], and are also referred to as the horizontal fiber system [15, Fig. 6b]. [^1^ These sources appear to consider CRE as part of LAL; this document treats CRE separately.] These neurons constitute a cholinergic pathway [1, Fig. 7C].

**Table IIX.**
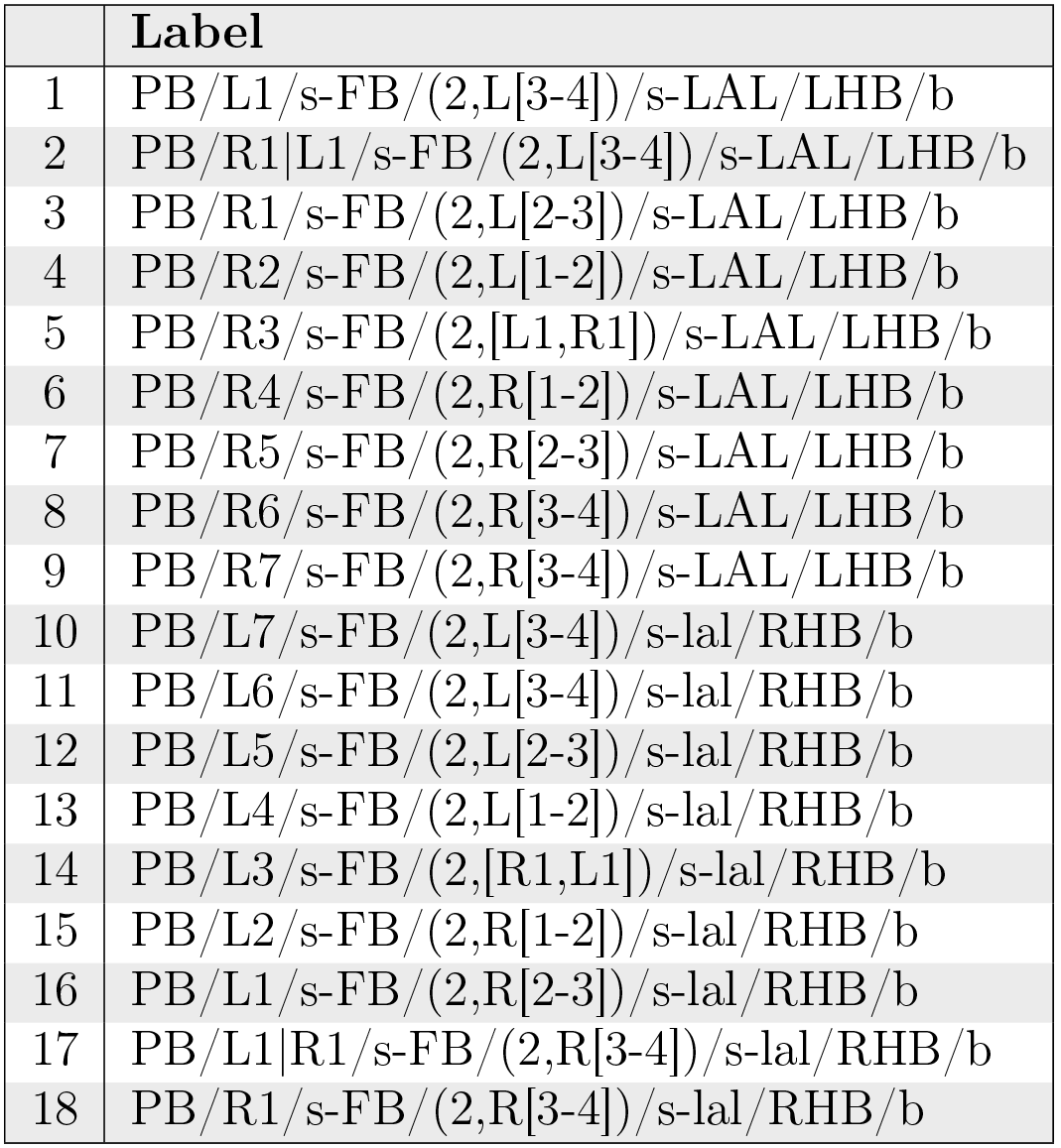
PB-FB-LAL neurons innervating layer 2 of FB[1, Fig. 6F].

**Fig. 21.**
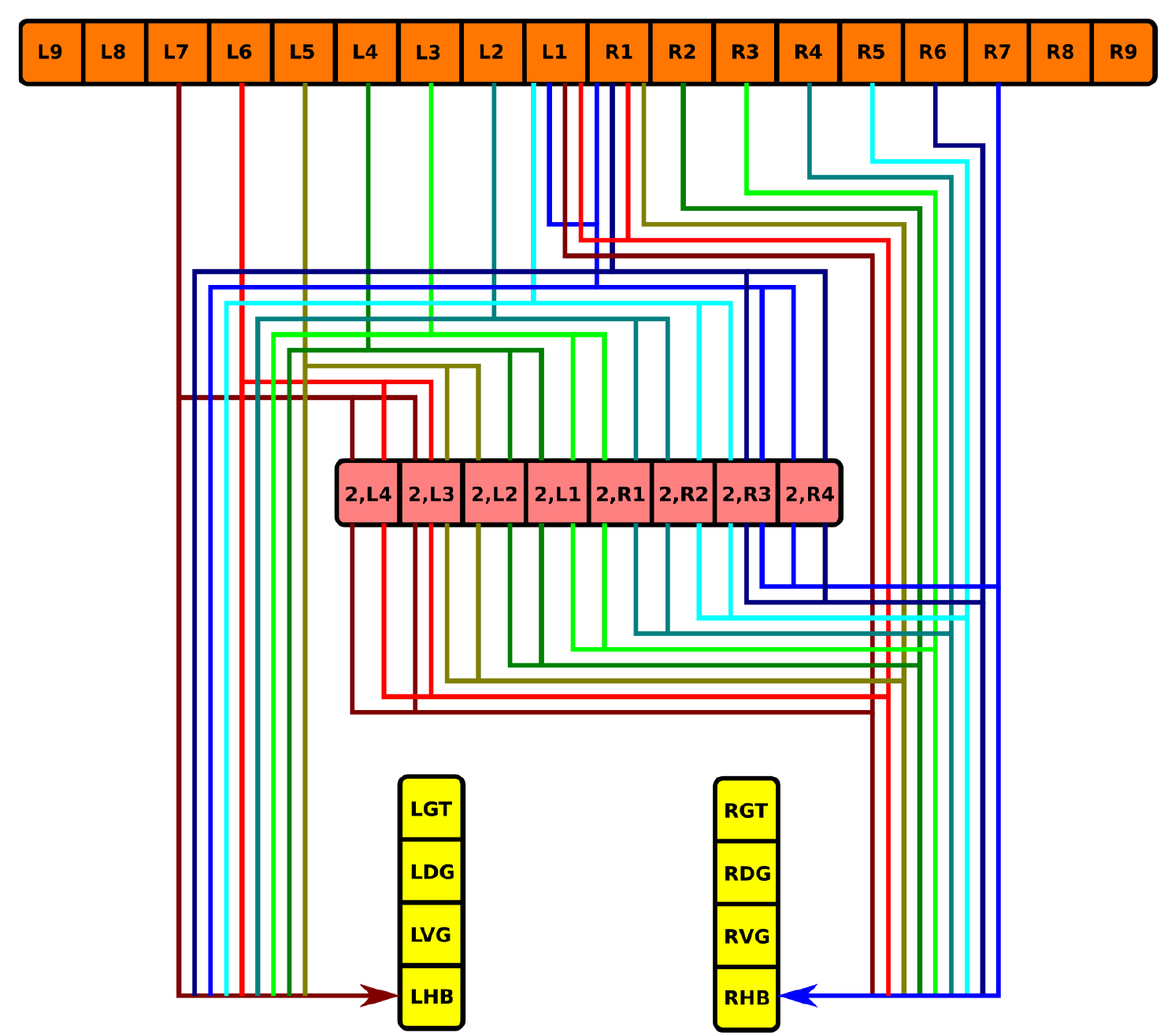
Neuropil innervation pattern for PB-FB-LAL neurons innervating layer 2 of FB (Table 19).

**Table XX.** PB-FB-LAL neurons [1, Fig. 6G].

|  | Label |
| --- | --- |
| 1 | PB/L1/s-FB/([1-4],L[3-4])/s-LAL/LHB/b |
| 2 | PB/L1\|R1/s-FB/([1-4],L[2-3])/s-LAL/LHB/b |
| 3 | PB/R1/s-FB/([1-4],L[1-2])/s-LAL/LHB/b |
| 4 | PB/R2/s-FB/([1-4],[L1,R1])/s-LAL/LHB/b |
| 5 | PB/R3/s-FB/([1-4],R[1-2])/s-LAL/LHB/b |
| 6 | PB/R4/s-FB/([1-4],R[2-3])/s-LAL/LHB/b |
| 7 | PB/R5/s-FB/([1-4],R[3-4])/s-LAL/LHB/b |
| 8 | PB/R6/s-FB/([1-4],R[3-4])/s-LAL/LHB/b |
| 9 | PB/L6/s-FB/([1-4],L[3-4])/s-lal/RHB/b |
| 10 | PB/L5/s-FB/([1-4],L[3-4])/s-lal/RHB/b |
| 11 | PB/L4/s-FB/([1-4],L[2-3])/s-lal/RHB/b |
| 12 | PB/L3/s-FB/([1-4],L[1-2])/s-lal/RHB/b |
| 13 | PB/L2/s-FB/([1-4],[L1,R1])/s-lal/RHB/b |
| 14 | PB/L1/s-FB/([1-4],R[1-2])/s-lal/RHB/b |
| 15 | PB/L1\|R1/s-FB/([1-4],R[2-3])/s-lal/RHB/b |
| 16 | PB/R1/s-FB/([1-4],R[3-4])/s-lal/RHB/b |

**Table XXI.**
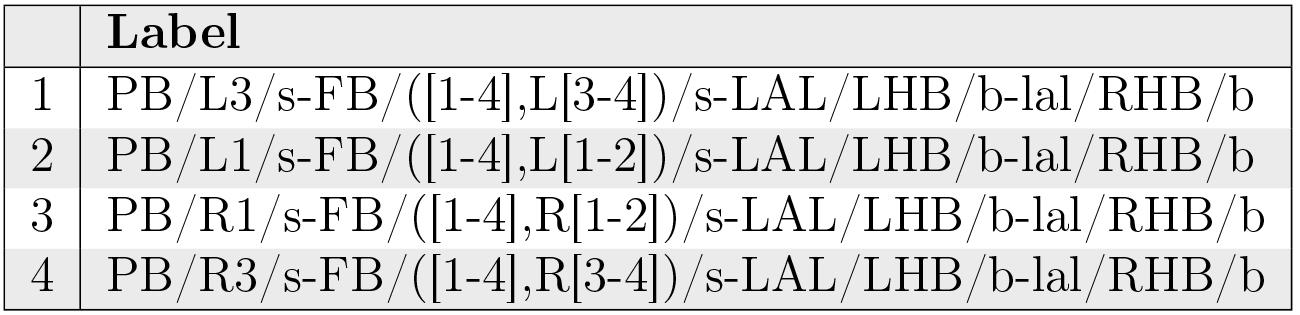
PB-FB-LAL neurons [1, Fig. 6G].

|  | Label |
| --- | --- |
| 1 | PB/L3/s-FB/([1-4],L[3-4])/s-LAL/LHB/b-lal/RHB/b |
| 2 | PB/L1/s-FB/([1-4],L[1-2])/s-LAL/LHB/b-lal/RHB/b |
| 3 | PB/R1/s-FB/([1-4],R[1-2])/s-LAL/LHB/b-lal/RHB/b |
| 4 | PB/R3/s-FB/([1-4],R[3-4])/s-LAL/LHB/b-lal/RHB/b |

**Fig. 22.**
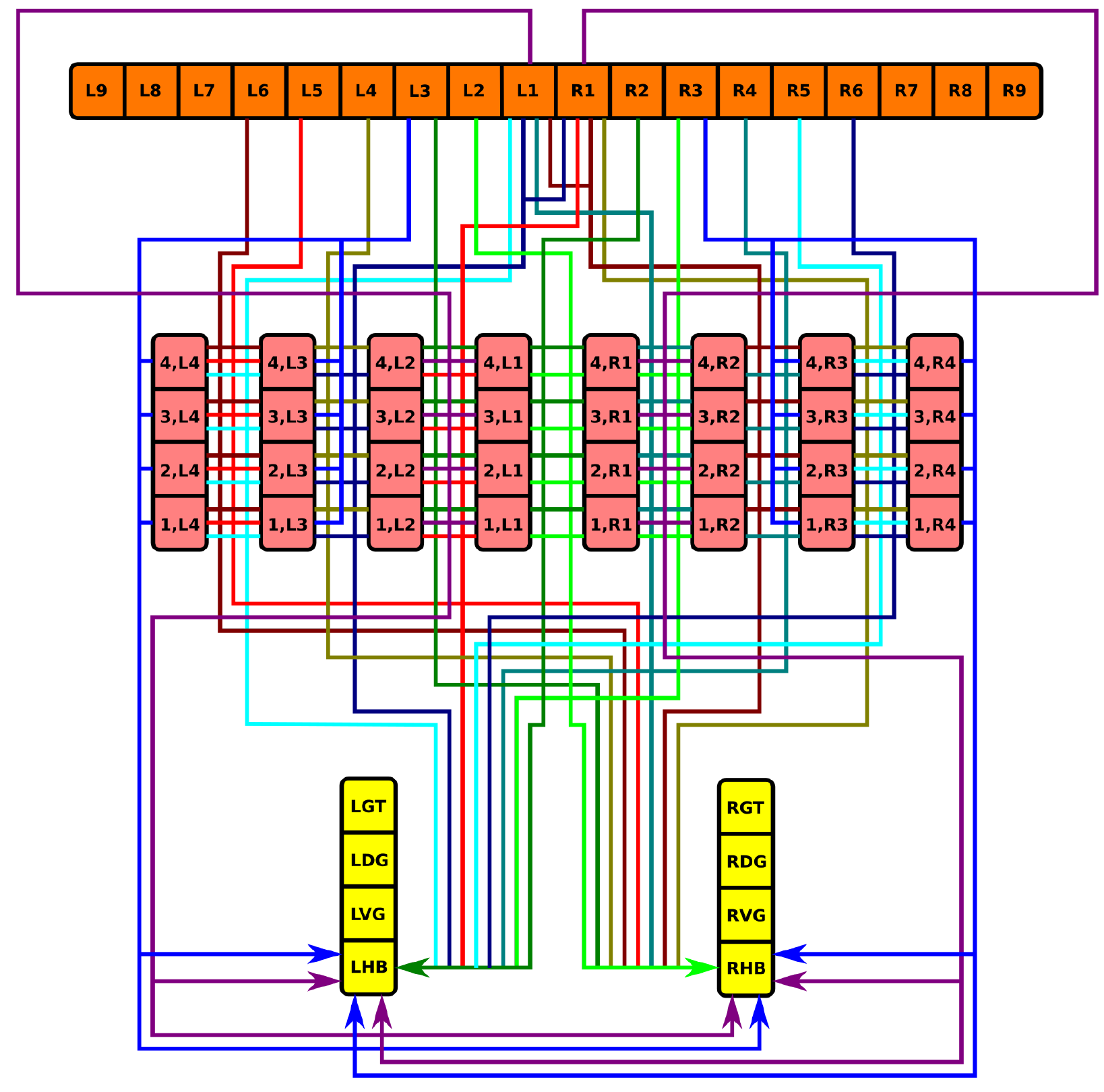
Neuropil innervation pattern for PB-FB-LAL neurons (Table 20, Table 21).

### 5.4.12 WED-PS-PB Projection Neurons

The neurons in Table 22 correspond to CCP-VMP-PB or CVP neurons in [1, Fig. 2] and receive input from the vision system. They constitute a cholinergic pathway [1, Fig. 7C].

**Table XXII.** WED-PS-PB neurons.

|  | Label |
| --- | --- |
| 1 | WED/L/s-PS/L/s-PB/L2\|L3/b |
| 2 | WED/L/s-PS/L/s-PB/L3\|L4/b |
| 3 | WED/L/s-PS/L/s-PB/L4\|L5/b |
| 4 | WED/L/s-PS/L/s-PB/L5\|L6/b |
| 5 | WED/L/s-PS/L/s-PB/L6\|L7/b |
| 6 | WED/L/s-PS/L/s-PB/L7\|L8/b |
| 7 | WED/L/s-PS/L/s-PB/L8\|L9/b |
| 8 | wed/R/s-ps/R/s-PB/L2\|L3/b |
| 9 | wed/R/s-ps/R/s-PB/L3\|L4/b |
| 10 | wed/R/s-ps/R/s-PB/L4\|L5/b |
| 11 | wed/R/s-ps/R/s-PB/L5\|L6/b |
| 12 | wed/R/s-ps/R/s-PB/L6\|L7/b |
| 13 | wed/R/s-ps/R/s-PB/L7\|L8/b |
| 14 | wed/R/s-ps/R/s-PB/L8\|L9/b |
| 15 | WED/L/s-PS/L/s-PB/R2\|R3/b |
| 16 | WED/L/s-PS/L/s-PB/R3\|R4/b |
| 17 | WED/L/s-PS/L/s-PB/R4\|R5/b |
| 18 | WED/L/s-PS/L/s-PB/R5\|R6/b |
| 19 | WED/L/s-PS/L/s-PB/R6\|R7/b |
| 20 | WED/L/s-PS/L/s-PB/R7\|R8/b |
| 21 | WED/L/s-PS/L/s-PB/R8\|R9/b |
| 22 | wed/R/s-ps/R/s-PB/R2\|R3/b |
| 23 | wed/R/s-ps/R/s-PB/R3\|R4/b |
| 24 | wed/R/s-ps/R/s-PB/R4\|R5/b |
| 25 | wed/R/s-ps/R/s-PB/R5\|R6/b |
| 26 | wed/R/s-ps/R/s-PB/R6\|R7/b |
| 27 | wed/R/s-ps/R/s-PB/R7\|R8/b |
| 28 | wed/R/s-ps/R/s-PB/R8\|R9/b |

**Fig. 23.**
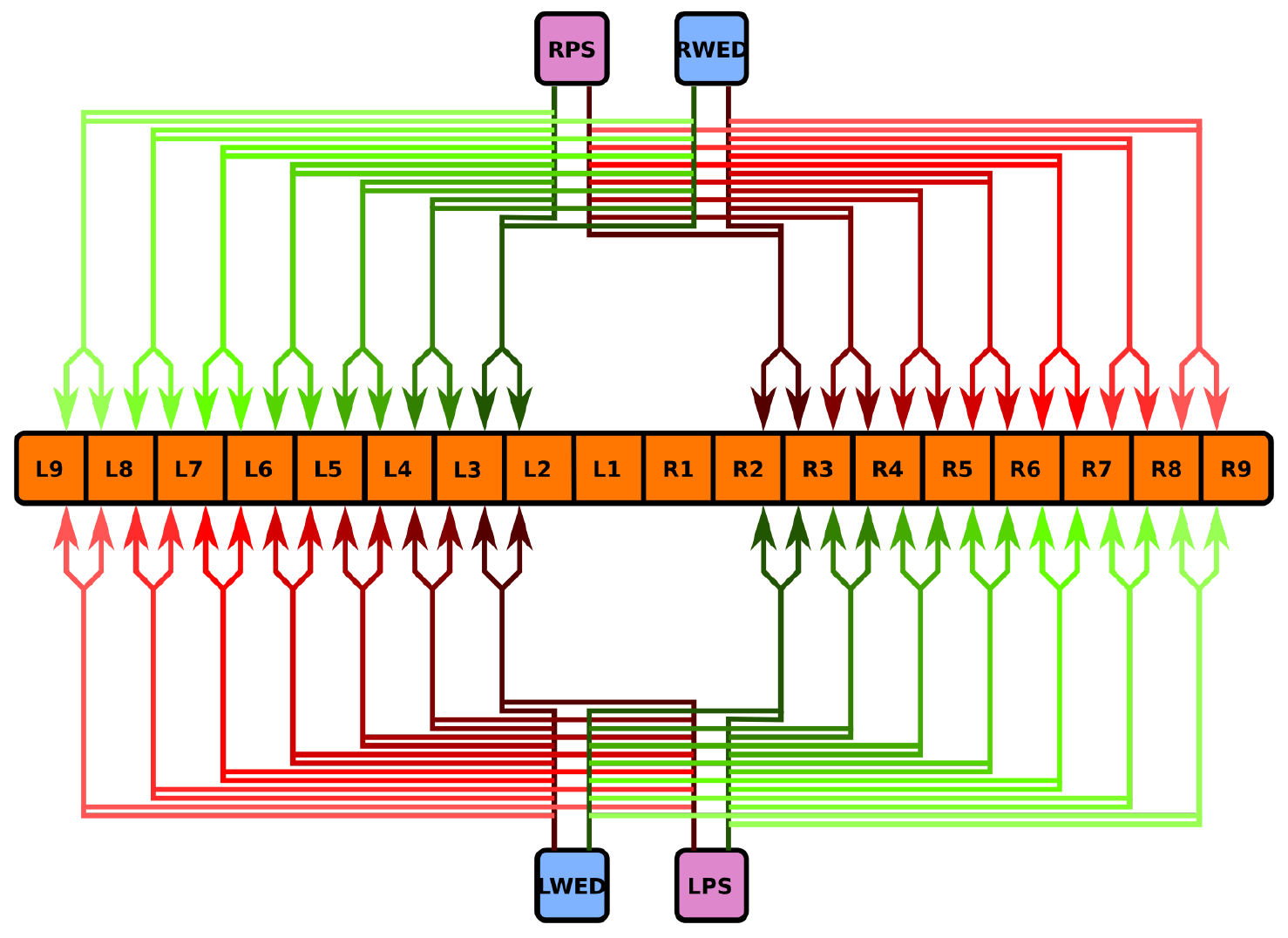
Neuropil innervation pattern for WED-PS-PB neurons (Table 22).

## 6 Generating an Executable Circuit Model

### 6.1 Neuron Organization

In light of the current lack of data regarding synapses between the various neurons identified in the central complex neuropils, data regarding the arborizations of these neurons was used to infer the presence or absence of synapses to generate an executable model of the central complex. Local and projection neurons were assigned to LPUs as indicated in Table 23.

**Table XXIII.**
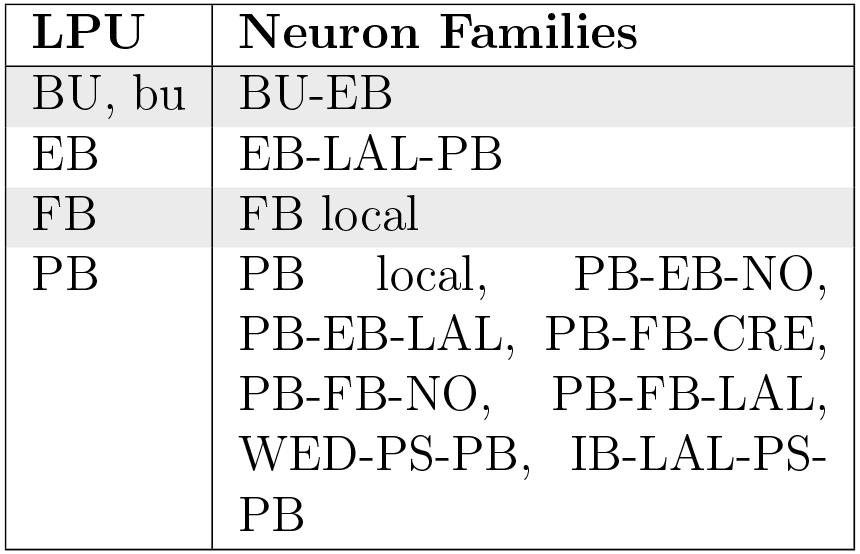
Assignment of neuron families to LPUs in generated CX model.

| LPU | Neuron Families |
| --- | --- |
| BU, bu | BU-EB |
| EB | EB-LAL-PB |
| FB | FB local |
| PB | PB local, PB-EB-NO, PB-EB-LAL, PB-FB-CRE, PB-FB-NO, PB-FB-LAL, WED-PS-PB, IB-LAL-PS-PB |

Although the BU-EB neurons have not been systematically characterized, available information regarding these neurons (§ 5.4.1) was used to hypothesize the following arborization structure for a total of 80 neurons in each hemisphere of the fly brain (Table 24).

**Table XXIV.** Hypothesized arborizations of BU-EB neurons. Each microglomerulus corresponds to a single neuron, where the “x” character in the neuron label is replaced with the microglomerulus number.

| Ring Neuron Name | Microglomerulus Range | Label |
| --- | --- | --- |
| R1 | 1..16 | BU/Lx/s-EB/(LR[1-8],[M,A],1)/b<br>bu/Rx/s-EB/(LR[1-8],[M,A],1)/b |
| R2 | 33..48 | BU/Lx/s-EB/(LR[1-8],A,[3,4])/b<br>bu/Rx/s-EB/(LR[1-8],A,[3,4])/b |
| R3 | 17..32 | BU/Lx/s-EB/(LR[1-8],A,[1,2])/b<br>bu/Rx/s-EB/(LR[1-8],A,[1,2])/b |
| R4m | 49..64 | BU/Lx/s-EB/(LR[1-8],A,4)/b<br>bu/Rx/s-EB/(LR[1-8],A,4)/b |
| R4d | 65..80 | BU/Lx/s-EB/(LR[1-8],M,4)/b<br>bu/Rx/s-EB/(LR[1-8],M,4)/b |

Likewise, we also hypothesized isomorphic sets of pontine neurons that link the following regions in FB based upon [15, p. 349]:

- nonadjacent segments in layers 1, 2, 4, and 5 (Table 25);
- adjacent segments within the same layer in layers 1-5 [15, Fig. 9a] (Table 26).
- adjacent layers within the same segment for layers 1-5 [15, Fig. 9a] (Table 27).
- nonadjacent layers within the same segment, with both presynaptic and postsynaptic terminals in each layer [15, Fig. 9b] (Table 28); based upon the latter source, we assume two sets of neurons connecting layers 1 and 8 and 2 and 7, respectively.

**Table XXV.** Hypothesized FB local neurons linking segments in layers 1, 2, 4, and 5, respectively.

|  | **Label** |
| --- | --- |
| 1 | FB/(1,L4)/s-FB/(1,R1)/b |
| 2 | FB/(1,L3)/s-FB/(1,R2)/b |
| 3 | FB/(1,L2)/s-FB/(1,R3)/b |
| 4 | FB/(1,L1)/s-FB/(1,R4)/b |
| 5 | FB/(1,R4)/s-FB/(1,L1)/b |
| 6 | FB/(1,R3)/s-FB/(1,L2)/b |
| 7 | FB/(1,R2)/s-FB/(1,L3)/b |
| 8 | FB/(1,R1)/s-FB/(1,L4)/b |
| 9 | FB/(2,L4)/s-FB/(2,R1)/b |
| 10 | FB/(2,L3)/s-FB/(2,R2)/b |
| 11 | FB/(2,L2)/s-FB/(2,R3)/b |
| 12 | FB/(2,L1)/s-FB/(2,R4)/b |
| 13 | FB/(2,R4)/s-FB/(2,L1)/b |
| 14 | FB/(2,R3)/s-FB/(2,L2)/b |
| 15 | FB/(2,R2)/s-FB/(2,L3)/b |
| 16 | FB/(2,R1)/s-FB/(2,L4)/b |
| 17 | FB/(4,L4)/s-FB/(4,R1)/b |
| 18 | FB/(4,L3)/s-FB/(4,R2)/b |
| 19 | FB/(4,L2)/s-FB/(4,R3)/b |
| 20 | FB/(4,L1)/s-FB/(4,R4)/b |
| 21 | FB/(4,R4)/s-FB/(4,L1)/b |
| 22 | FB/(4,R3)/s-FB/(4,L2)/b |
| 23 | FB/(4,R2)/s-FB/(4,L3)/b |
| 24 | FB/(4,R1)/s-FB/(4,L4)/b |
| 25 | FB/(5,L4)/s-FB/(5,R1)/b |
| 26 | FB/(5,L3)/s-FB/(5,R2)/b |
| 27 | FB/(5,L2)/s-FB/(5,R3)/b |
| 28 | FB/(5,L1)/s-FB/(5,R4)/b |
| 29 | FB/(5,R4)/s-FB/(5,L1)/b |
| 30 | FB/(5,R3)/s-FB/(5,L2)/b |
| 31 | FB/(5,R2)/s-FB/(5,L3)/b |
| 32 | FB/(5,R1)/s-FB/(5,L4)/b |

**Table XXVI.** Hypothesized FB local neurons linking adjacent segments within the same layer in layers 1-5.

|  | Label |
| --- | --- |
| 1 | FB/(1,L4)/s-FB/(1,L3)/b |
| 2 | FB/(1,L3)/s-FB/(1,L2)/b |
| 3 | FB/(1,L2)/s-FB/(1,L1)/b |
| 4 | FB/(1,L1)/s-FB/(1,R1)/b |
| 5 | FB/(1,R1)/s-FB/(1,R2)/b |
| 6 | FB/(1,R2)/s-FB/(1,R3)/b |
| 7 | FB/(1,R3)/s-FB/(1,R4)/b |
| 8 | FB/(2,L4)/s-FB/(2,L3)/b |
| 9 | FB/(2,L3)/s-FB/(2,L2)/b |
| 10 | FB/(2,L2)/s-FB/(2,L1)/b |
| 11 | FB/(2,L1)/s-FB/(2,R1)/b |
| 12 | FB/(2,R1)/s-FB/(2,R2)/b |
| 13 | FB/(2,R2)/s-FB/(2,R3)/b |
| 14 | FB/(2,R3)/s-FB/(2,R4)/b |
| 15 | FB/(3,L4)/s-FB/(3,L3)/b |
| 16 | FB/(3,L3)/s-FB/(3,L2)/b |
| 17 | FB/(3,L2)/s-FB/(3,L1)/b |
| 18 | FB/(3,L1)/s-FB/(3,R1)/b |
| 19 | FB/(3,R1)/s-FB/(3,R2)/b |
| 20 | FB/(3,R2)/s-FB/(3,R3)/b |
| 21 | FB/(3,R3)/s-FB/(3,R4)/b |
| 22 | FB/(4,L4)/s-FB/(4,L3)/b |
| 23 | FB/(4,L3)/s-FB/(4,L2)/b |
| 24 | FB/(4,L2)/s-FB/(4,L1)/b |
| 25 | FB/(4,L1)/s-FB/(4,R1)/b |
| 26 | FB/(4,R1)/s-FB/(4,R2)/b |
| 27 | FB/(4,R2)/s-FB/(4,R3)/b |
| 28 | FB/(4,R3)/s-FB/(4,R4)/b |
| 29 | FB/(5,L4)/s-FB/(5,L3)/b |
| 30 | FB/(5,L3)/s-FB/(5,L2)/b |
| 31 | FB/(5,L2)/s-FB/(5,L1)/b |
| 32 | FB/(5,L1)/s-FB/(5,R1)/b |
| 33 | FB/(5,R1)/s-FB/(5,R2)/b |
| 34 | FB/(5,R2)/s-FB/(5,R3)/b |
| 35 | FB/(5,R3)/s-FB/(5,R4)/b |

**Table XXVII.** Hypothesized FB local neurons linking adjacent layers within the same segment for layers 1-5.

|  | **Label** |
| --- | --- |
| 1 | FB/(5,L1)/s-FB/(4,L1)/b |
| 2 | FB/(4,L1)/s-FB/(3,L1)/b |
| 3 | FB/(3,L1)/s-FB/(2,L1)/b |
| 4 | FB/(2,L1)/s-FB/(1,L1)/b |
| 5 | FB/(5,L2)/s-FB/(4,L2)/b |
| 6 | FB/(4,L2)/s-FB/(3,L2)/b |
| 7 | FB/(3,L2)/s-FB/(2,L2)/b |
| 8 | FB/(2,L2)/s-FB/(1,L2)/b |
| 9 | FB/(5,L3)/s-FB/(4,L3)/b |
| 10 | FB/(4,L3)/s-FB/(3,L3)/b |
| 11 | FB/(3,L3)/s-FB/(2,L3)/b |
| 12 | FB/(2,L3)/s-FB/(1,L3)/b |
| 13 | FB/(5,L4)/s-FB/(4,L4)/b |
| 14 | FB/(4,L4)/s-FB/(3,L4)/b |
| 15 | FB/(3,L4)/s-FB/(2,L4)/b |
| 16 | FB/(2,L4)/s-FB/(1,L4)/b |
| 17 | FB/(5,R1)/s-FB/(4,R1)/b |
| 18 | FB/(4,R1)/s-FB/(3,R1)/b |
| 19 | FB/(3,R1)/s-FB/(2,R1)/b |
| 20 | FB/(2,R1)/s-FB/(1,R1)/b |
| 21 | FB/(5,R2)/s-FB/(4,R2)/b |
| 22 | FB/(4,R2)/s-FB/(3,R2)/b |
| 23 | FB/(3,R2)/s-FB/(2,R2)/b |
| 24 | FB/(2,R2)/s-FB/(1,R2)/b |
| 25 | FB/(5,R3)/s-FB/(4,R3)/b |
| 26 | FB/(4,R3)/s-FB/(3,R3)/b |
| 27 | FB/(3,R3)/s-FB/(2,R3)/b |
| 28 | FB/(2,R3)/s-FB/(1,R3)/b |
| 29 | FB/(5,R4)/s-FB/(4,R4)/b |
| 30 | FB/(4,R4)/s-FB/(3,R4)/b |
| 31 | FB/(3,R4)/s-FB/(2,R4)/b |
| 32 | FB/(2,R4)/s-FB/(1,R4)/b |

**Table XXVIII.** Hypothesized FB local neurons linking nonadjacent layers within the same segment.

|  | Label |
| --- | --- |
| 1 | FB/(1,L1)/sb-FB/(8,L1)/sb |
| 2 | FB/(1,L2)/sb-FB/(8,L2)/sb |
| 3 | FB/(1,L3)/sb-FB/(8,L3)/sb |
| 4 | FB/(1,L4)/sb-FB/(8,L4)/sb |
| 5 | FB/(1,R1)/sb-FB/(8,R1)/sb |
| 6 | FB/(1,R2)/sb-FB/(8,R2)/sb |
| 7 | FB/(1,R3)/sb-FB/(8,R3)/sb |
| 8 | FB/(1,R4)/sb-FB/(8,R4)/sb |
| 9 | FB/(2,L1)/sb-FB/(7,L1)/sb |
| 10 | FB/(2,L2)/sb-FB/(7,L2)/sb |
| 11 | FB/(2,L3)/sb-FB/(7,L3)/sb |
| 12 | FB/(2,L4)/sb-FB/(7,L4)/sb |
| 13 | FB/(2,R1)/sb-FB/(7,R1)/sb |
| 14 | FB/(2,R2)/sb-FB/(7,R2)/sb |
| 15 | FB/(2,R3)/sb-FB/(7,R3)/sb |
| 16 | FB/(2,R4)/sb-FB/(7,R4)/sb |

### 6.2 Executable Circuit Generation

To infer the presence of synaptic connections between neurons, the above neuron names were loaded into a NeuroArch database [36] in accordance with its data model. Using a parser for the grammar described in § 2.2, each neuron’s name was parsed to extract its constituent arborization records (Table 29); these records were reinserted into the database as ArborizationData nodes and connected to the Neuron nodes created for the neuron in each family listed above.

**Table XXIX.** Fields in ArborizationData node. Region strings or tuples conform to the formats described in § 3.

| Field | Data Type | Sample Values |
| --- | --- | --- |
| neurite | set of ‘b’ or ‘s’ | [b], [b, s] |
| neuropil | string | PB, EB |
| region | set of strings or tuples | [L1], [(1, R1)] |

After extraction of arborization data, all pairs of neurons in the database were compared to find those pairs with geometrically overlapping arborizations and differing neurite types (i.e. presynaptic versus postsynaptic). This resulted in the creation of Synapse nodes that were connected to the associated Neuron node pairs in NeuroArch’s database.

To illustrate the synapse inference algorithm’s operation, consider the neurons EB/ ([R3,R5],[P,M],[1–4])/s-EB/(R4,[P,M],[1–4])/b-LAL/RDG/b-PB/L3/b (Table 9) and PB/L4/s-EB/2/b-LAL/RVG/b (Table 12). Since the region EB/(R3,P,[1-4])/s overlaps with region EB/2/b (Table 1) and the terminal types of the two neurons in the overlapping region differ, we infer the presence of a synapse with information flow from the latter neuron to the former. Figs. 24, 25, 26, 27, 28, 29, 30, 31, and 32 respectively depict the inferred synapses between PB local neurons (Table 7), between PB local and PB-EB-LAL neurons (Table 12), between PB local and PB-EB-NO neurons (Table 11), between EB-LAL-PB neurons (Table 9), between EB-LAL-PB and PB local neurons, between EB-LAL-PB and PB-EB-LAL neurons, between EB-LAL-PB and PB-EB-NO neurons, between PB-EB-LAL and EB-LAL-PB neurons, and between PB-EB-NO and EB-LAL-PB neurons; yellow nodes correspond to PB local neurons, pink nodes correspond to PB-EB-LAL neurons, orange nodes correspond to PB-EB-NO neurons, and green nodes correspond to EB-LAL-PB neurons. In all of these figures, each arrow head corresponds to a single inferred synapse. The neuron numbers in the figures correspond to those in the respective tables of neuron names. In all of the following connectivity matrices, a black square denotes the presence of a connection linking a presynaptic neuron on the the y-axis to a postsynaptic neuron on the x-axis.

**Fig. 24.**
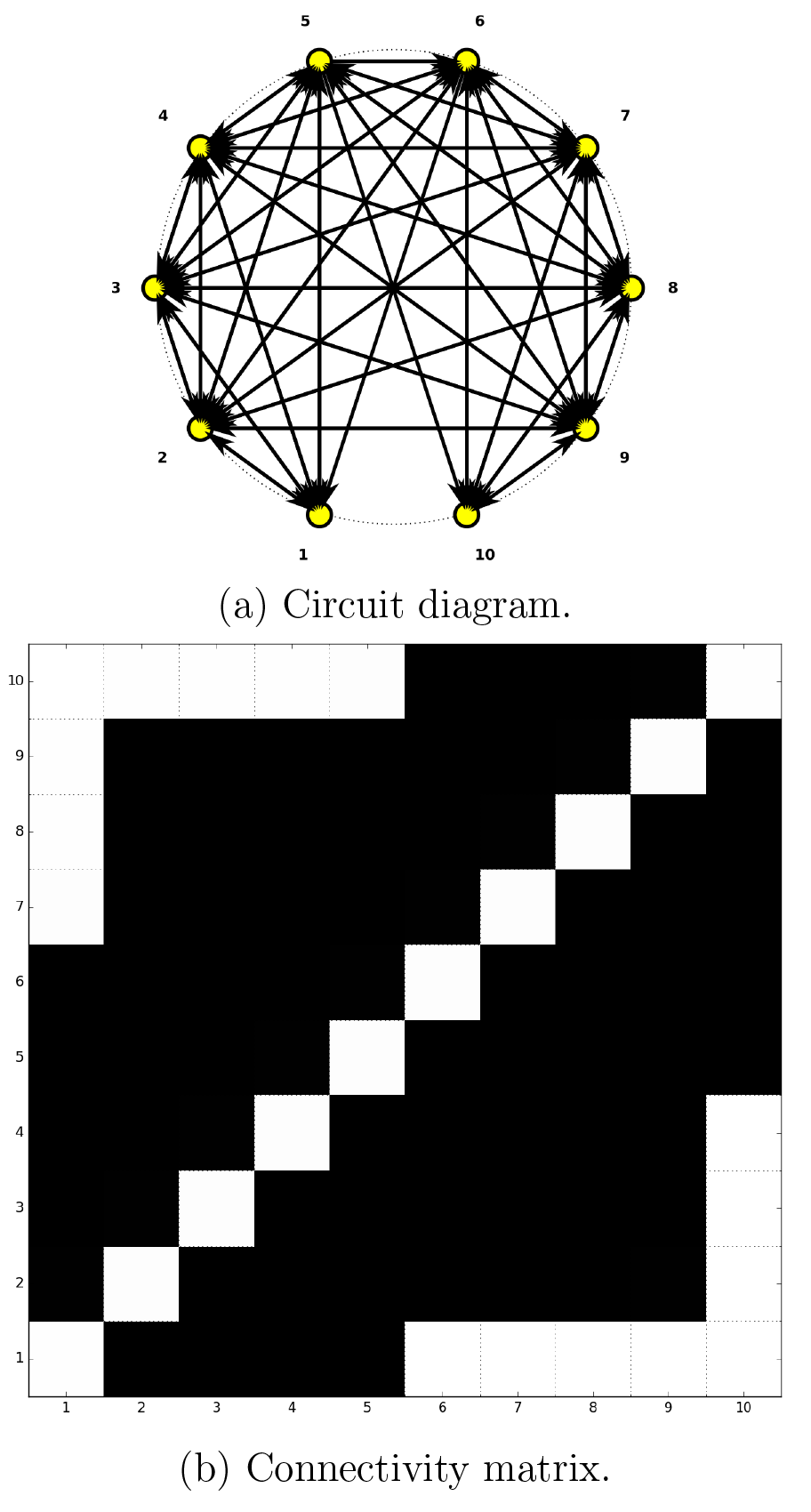
Inferred synapses between PB local neurons (Table 7).

**Fig. 25.**
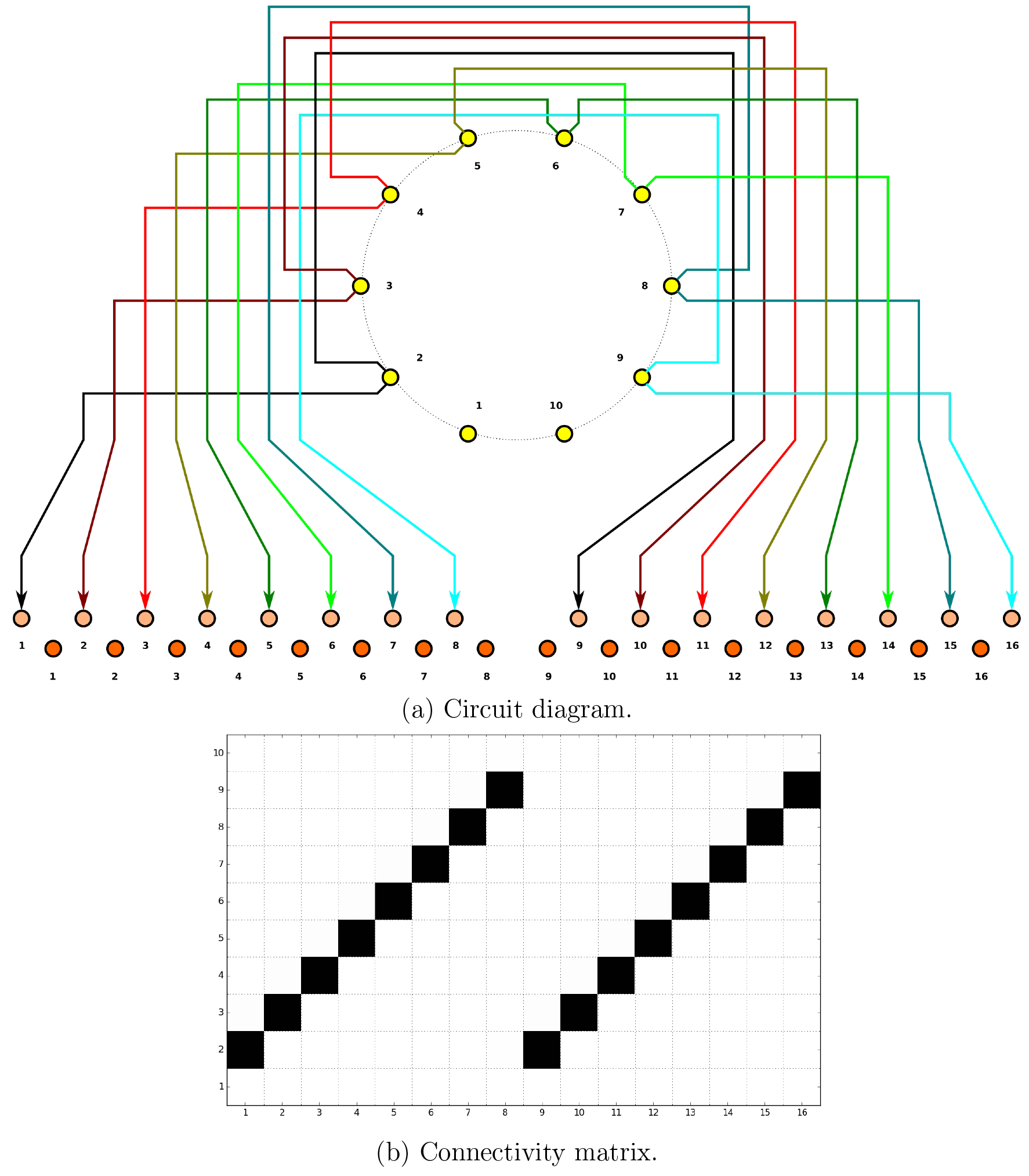
Inferred synapses between PB local (Table 7) and PB-EB-LAL (Table 12) neurons.

**Fig. 26.**
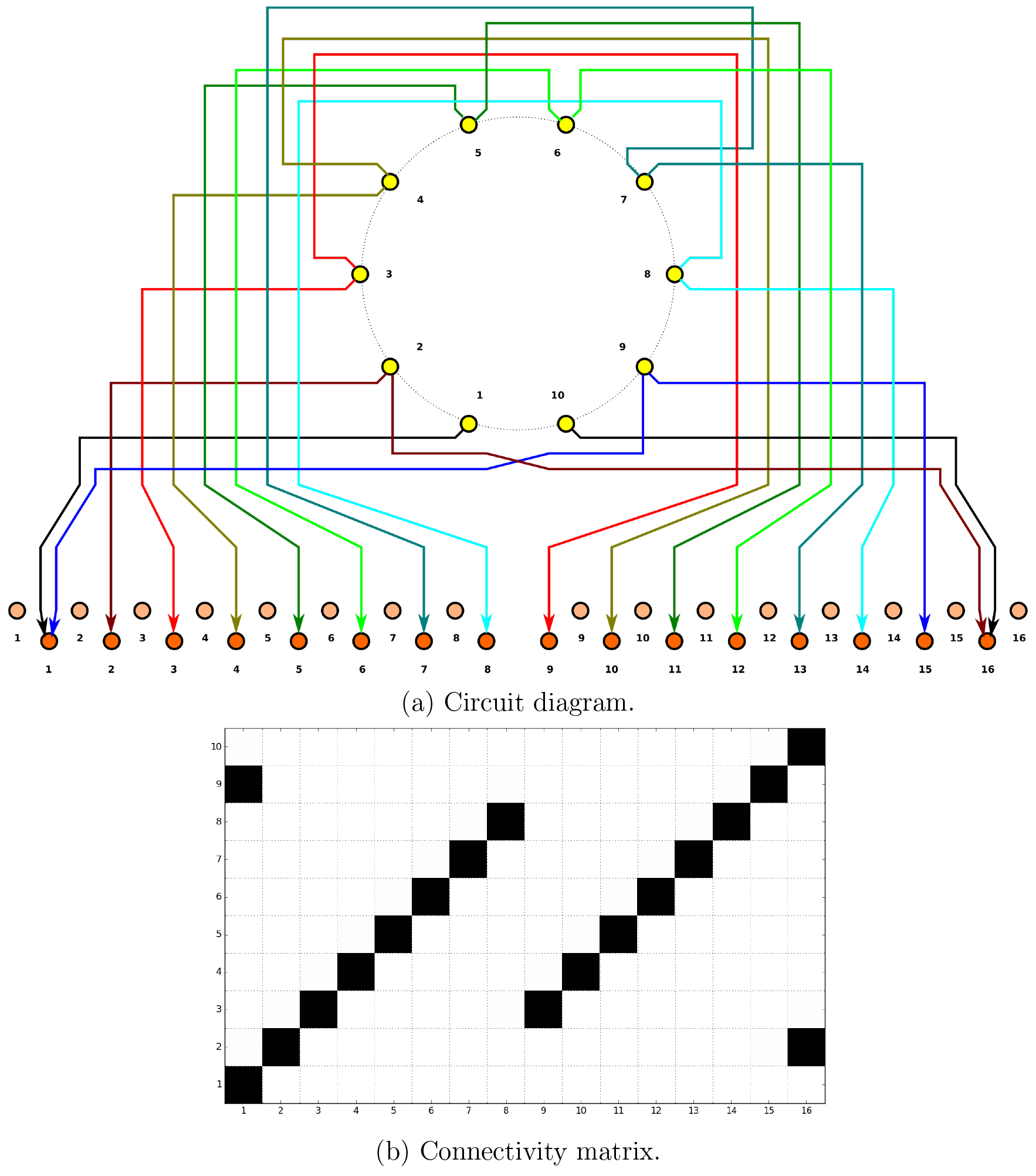
Inferred synapses between PB local (Table 7) and PB-EB-NO (Table 11) neurons.

**Fig. 27.**
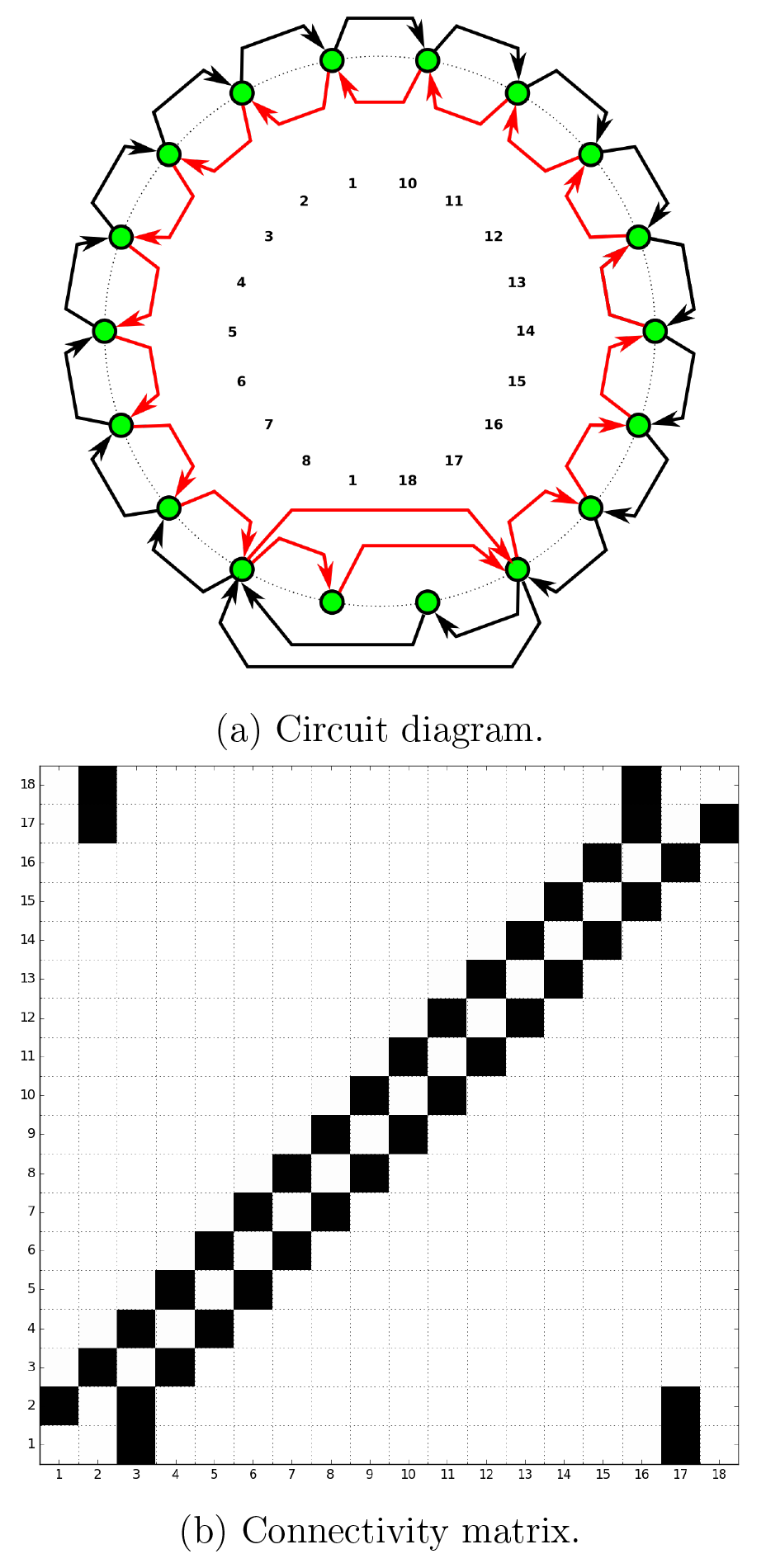
Inferred synapses between EB-LAL-PB (Table 9) neurons.

**Fig. 28.**
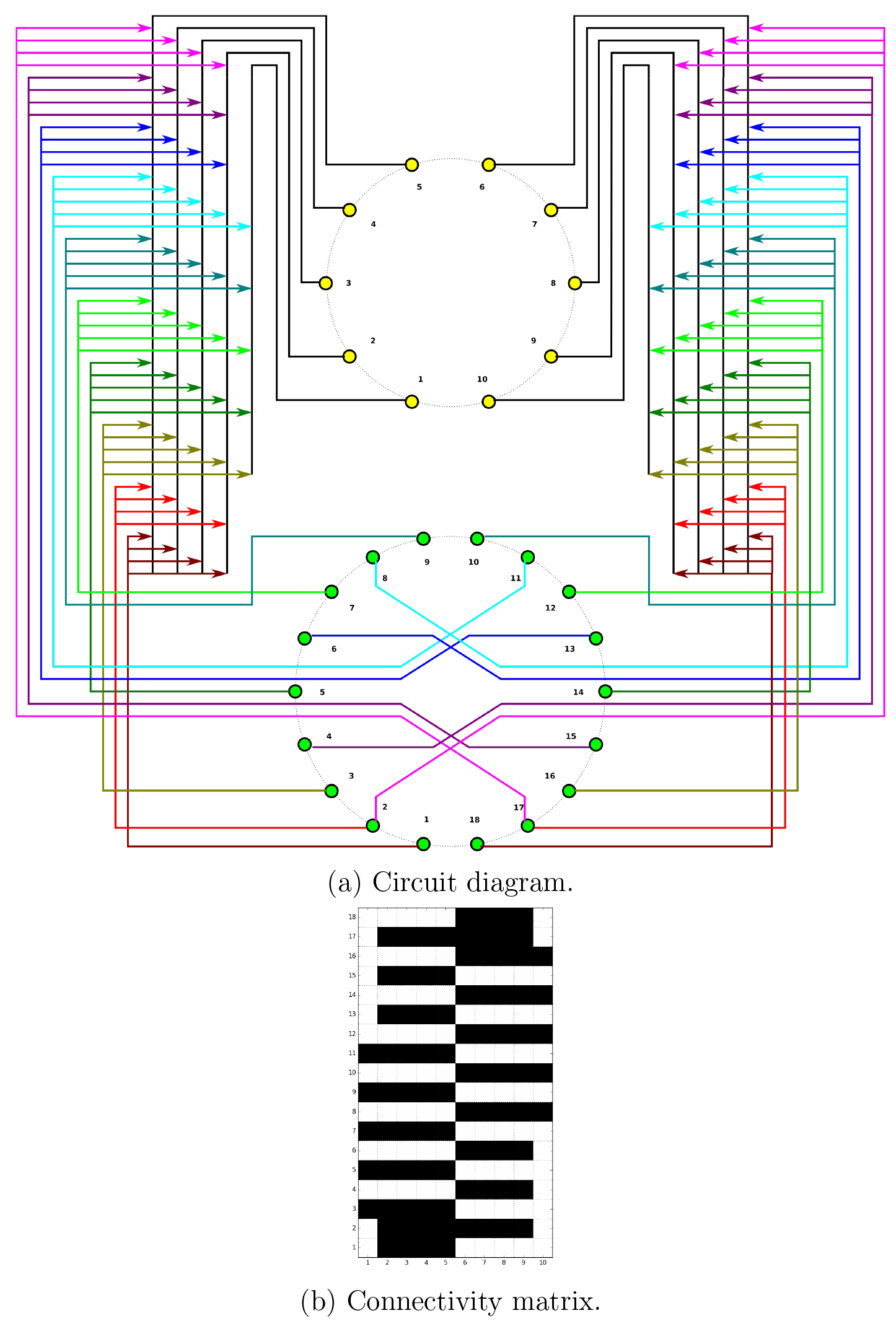
Inferred synapses between EB-LAL-PB (Table 9) and PB local (Table 7) neurons.

**Fig. 29.**
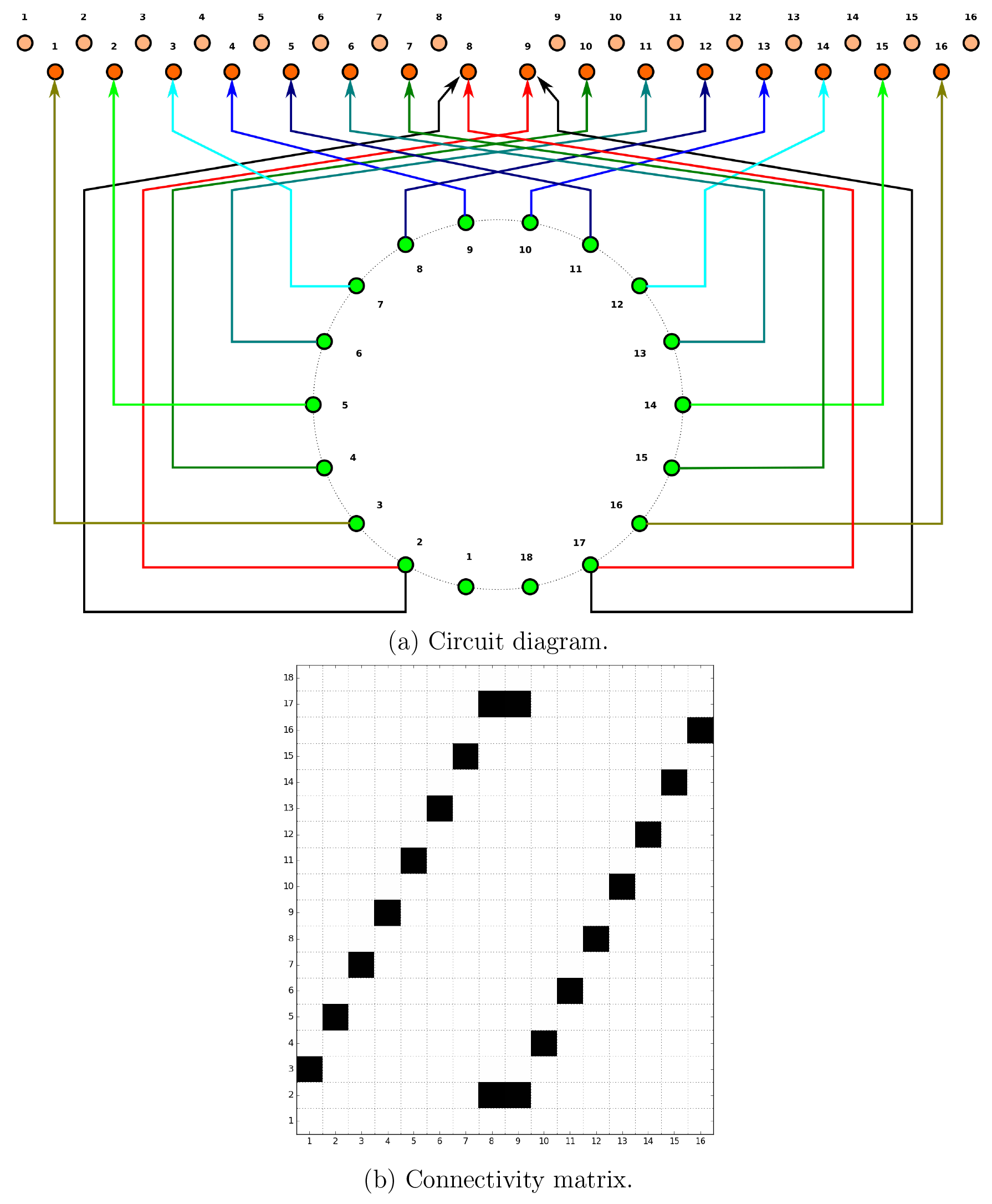
Inferred synapses between EB-LAL-PB (Table 9) and PB-EB-LAL (Table 12)
neurons.

**Fig. 30.**
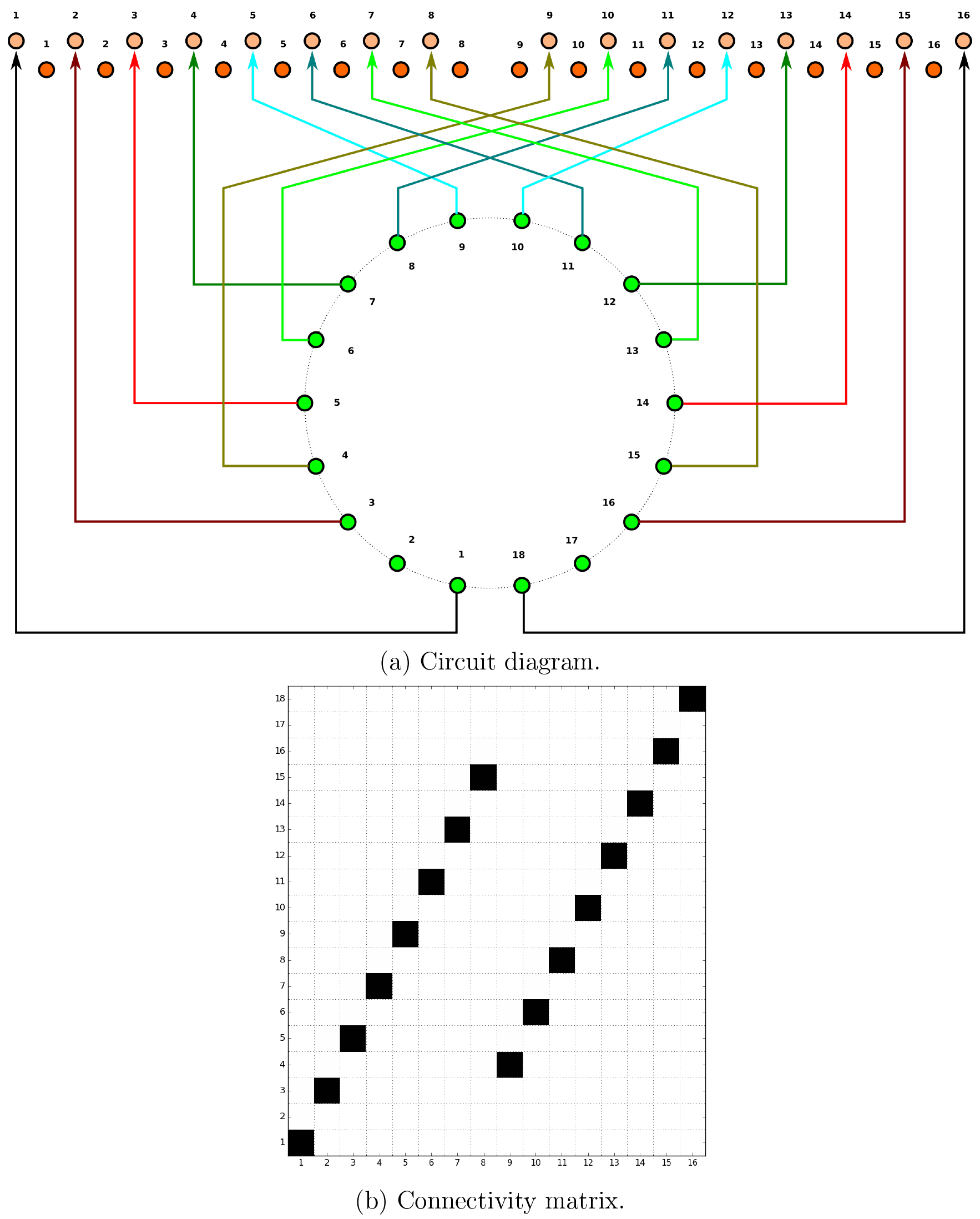
Inferred synapses between EB-LAL-PB (Table 9) and PB-EB-NO (Table 11) neurons.

**Fig. 31.**
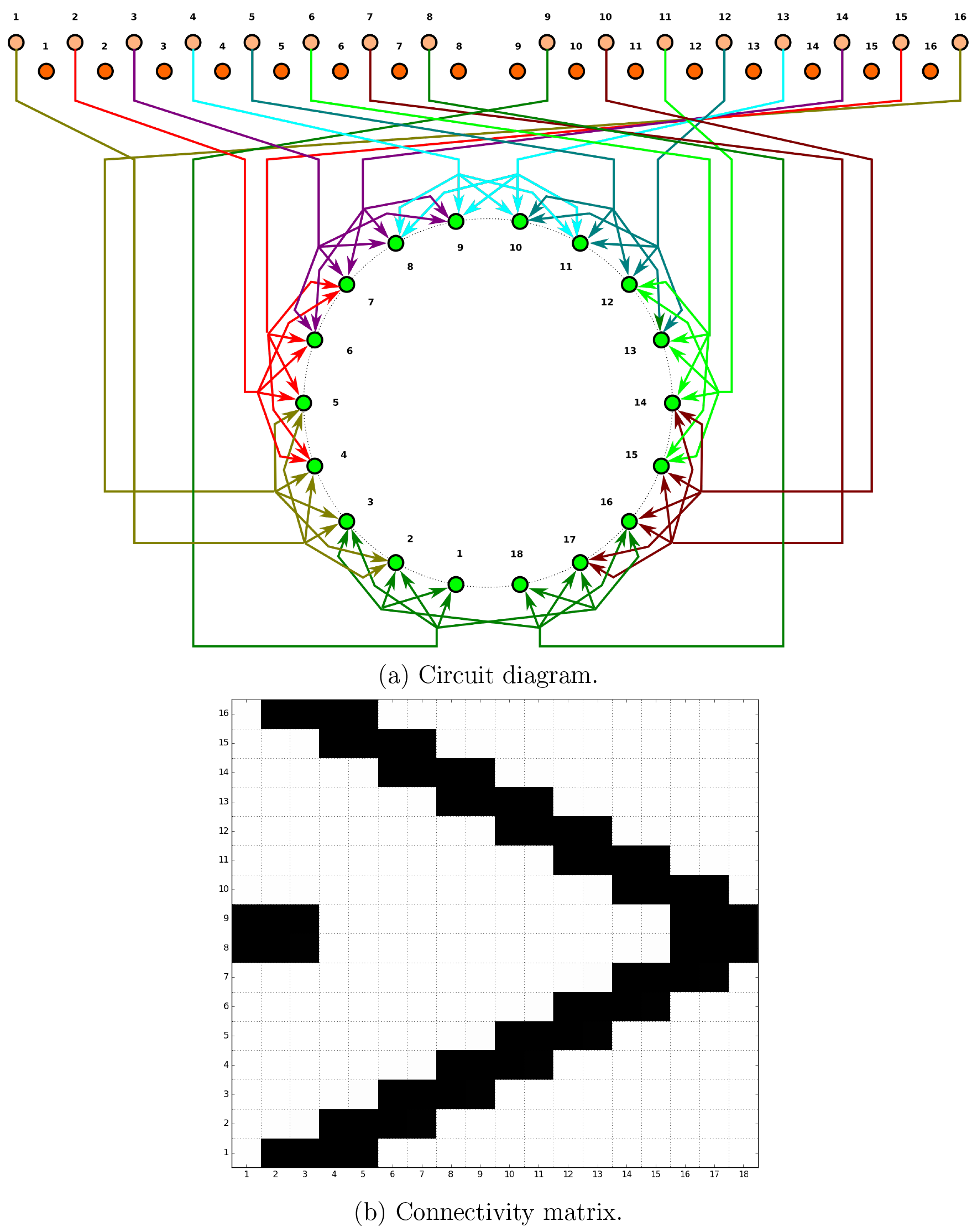
Inferred synapses between PB-EB-LAL (Table 12) and EB-LAL-PB (Table 9) neurons.

**Fig. 32.**
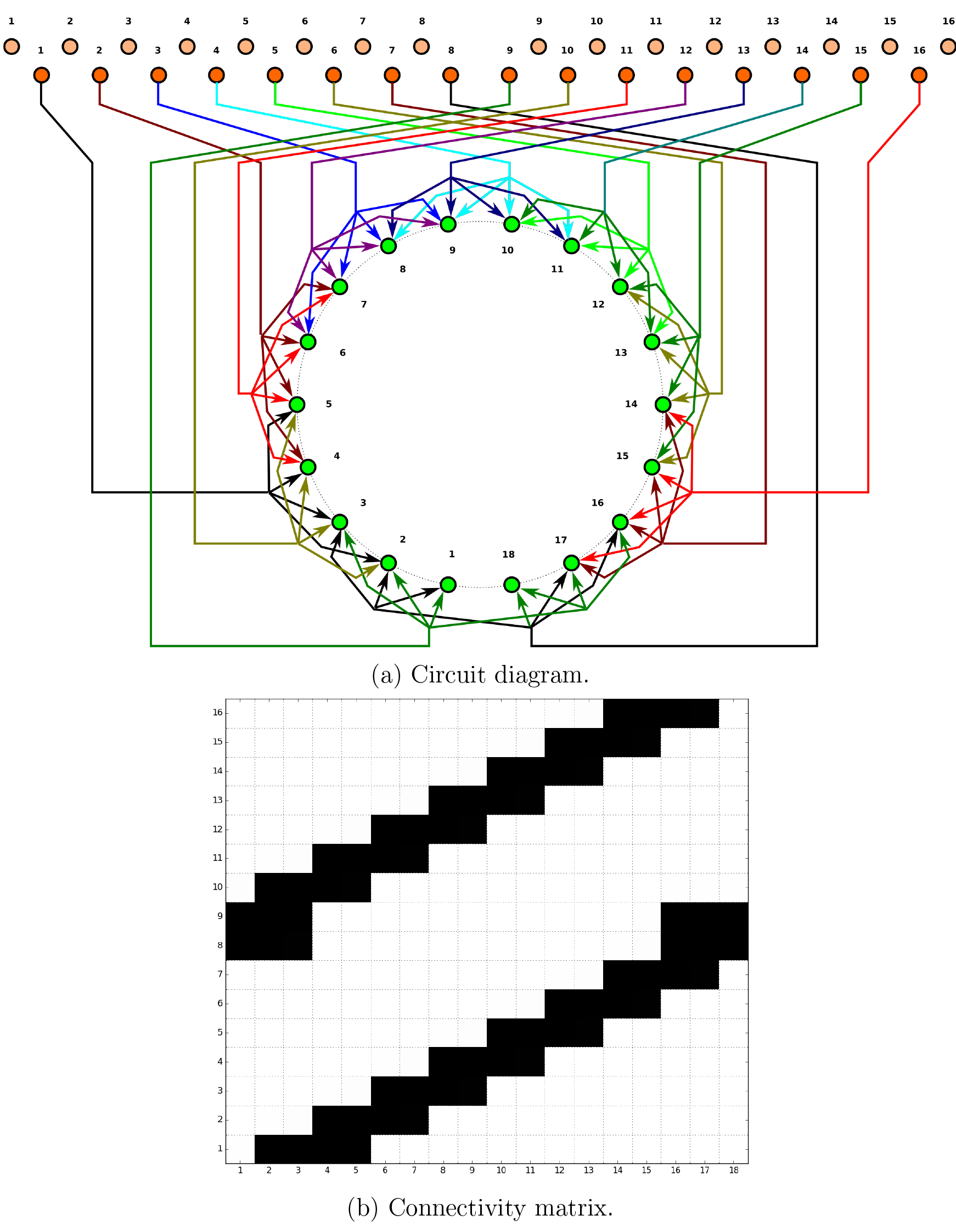
Inferred synapses between PB-EB-NO (Table 11) and EB-LAL-PB (Table 9) neurons.

Neuron and synapse models were instantiated for each respective biological neuron and synapse in NeuroArch’s database to construct LPUs corresponding to the BU, FB, EB, and PB neuropils. All neurons were modeled as Leaky Integrate-and-Fire neurons, and all synapses modeled to produce alpha function responses to presynaptic spikes. Communication ports were created for every neuron model instance comprised by one LPU connected to a synapse model instance in the other neuropil; the connectivity pattern linking the ports associated with the neuron and synapse models was also added to the NeuroArch database. NeuroArch’s API was used to extract the constructed LPUs and patterns and dispatch them to Neurokernel [37] for execution.

### 6.3 Executing the Circuit

To test the executability of the generated circuit and its ability to respond to input data, the generated model was driven by a simple visual stimulus consisting of an illuminated vertical bar proceeding horizontally across the 2D visual space (Fig. 33). Since the central complex neuropils do not receive direct connections from the vision neuropils, the visual stimulus was passed into three banks of receptive fields whose outputs were respectively provided to BU, bu, and PB as input (Fig. 34). The receptive fields for BU and bu each consisted of 80 evenly spaced 2D grids of circular Gaussians that correspond to one of the microglomeruli in BU; each receptive field was connected to one BU-EB neuron such that the 16 neurons in each of the 5 groups described in § 5.4.1 processed input from a rectangle occupying 5 of the 2D visual space. The receptive fields for PB consisted of 18 vertical rectangular regions with a constant magnitude; each receptive field was connected to all local and projection neurons that innervated the glomerulus corresponding to the receptive field region. The responses of the neurons in each family to the two input signals are organized in the same order in the respective raster plots.

**Fig. 33.**
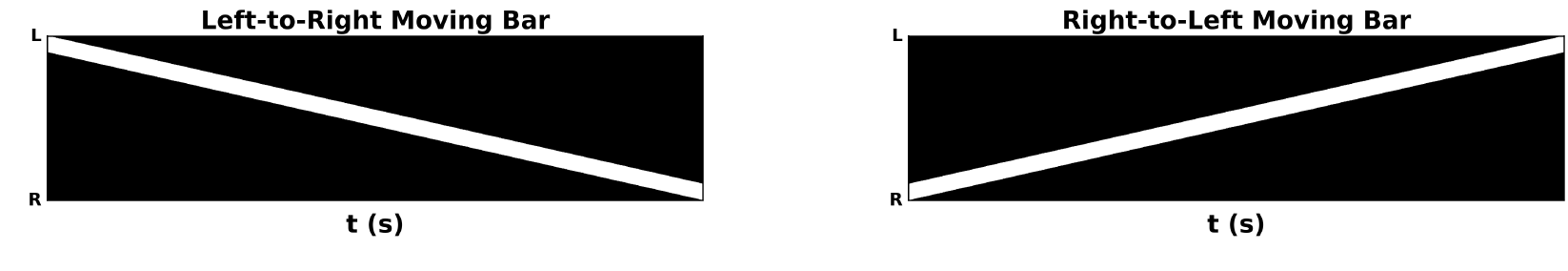
Moving bar visual input to generated CX model. The plots depict the movement of an illuminated vertical bar horizontally across a dark background.

**Fig. 34.**
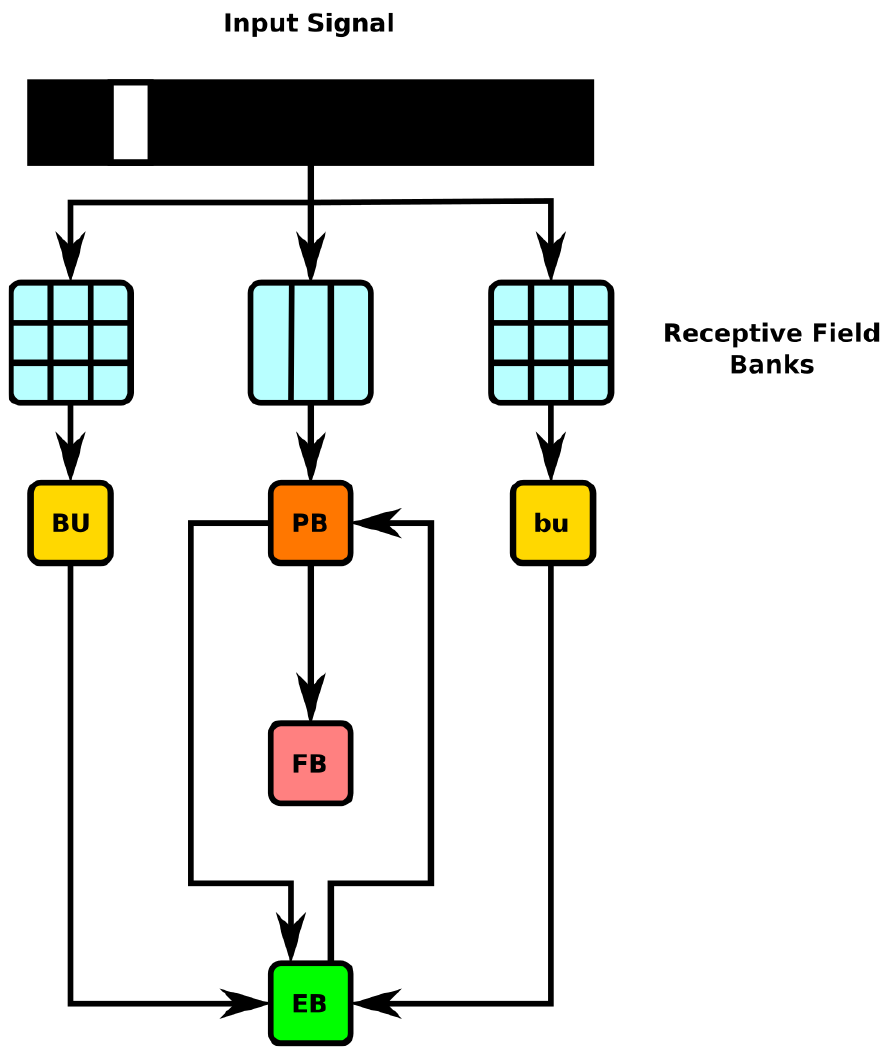
Schematic of information flow in generated CX model. 2D visual signals are passed through rectangular grids of Gaussian receptive fields whose outputs drive BU-EB neurons and through a bank of vertical rectangular receptive fields whose outputs drive neurons that innervate the PB glomeruli. The generated model only comprises neurons that innervate the depicted LPUs (BU, bu, EB, FB, and PB).

### 6.4 Use Cases

The NeuroArch and Neurokernel pipeline used to generate the CX model described above enables analysis and manipulation of the model using computational analogues to experimental techniques.

#### 6.4.1 Virtual Electrophysiology

One can use NeuroArch/Neurokernel to concurrently probe the responses different sets of neurons in multiple neuropils in a computational experiment. Figs. 35 and 36 depict the responses of neurons innervating the PB and BU/bu neuropils to the signal depicted in Fig. 33.

**Fig. 35.**
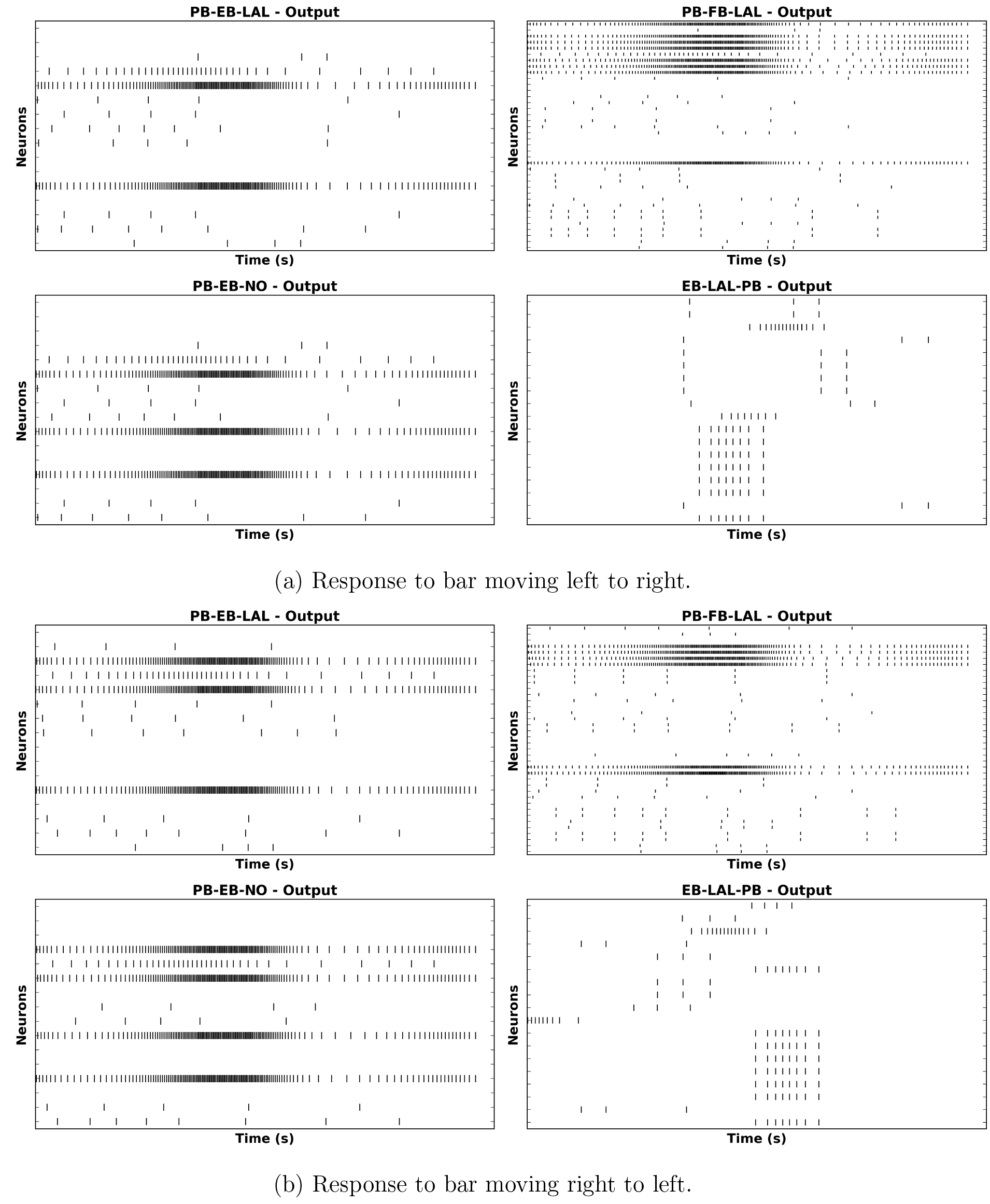
Response of CX projection neurons innervating PB to moving bar input.

**Fig. 36.**
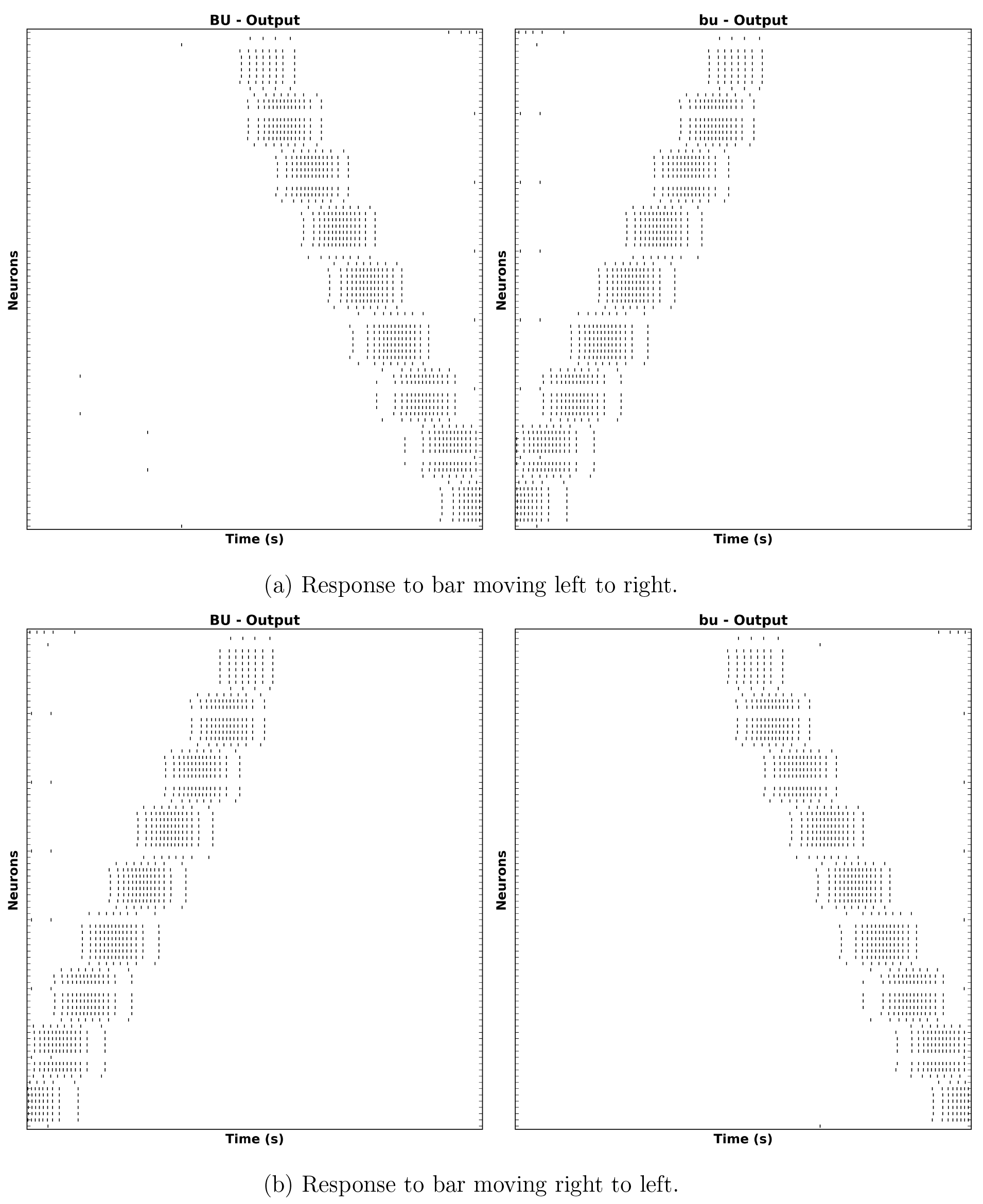
Response of CX projection neurons innervating BU/bu to moving bar input.

#### 6.4.2 Virtual Genetic Manipulation

To test hypotheses regarding incompletely characterized parts of the fly brain, one can create models that either attempt to replicate abnormal behaviors or emulate abnormal circuit structures observed in different mutant fly strains. For example, one can attempt to model phenotypes corresponding to mutations affecting the structure of PB (e.g., *no bridge, tay bridge,* etc.) by altering the PB model generation process accordingly. Given that these mutations are known to alter the fly’s step length [4, p. 7] and since neurons innervating the motor ganglia are known to be postsynaptic to those that innervate LAL, it is reasonable to expect that analogous modifications to the structure of PB may alter the observed output of CX projection neurons that innervate LAL.

We used NeuroArch to emulate the no *bridge* mutant by altering the PB local neurons to remove all local connections between the left and right sides of PB and positing the existence of additional local neurons caused by the mutation (Fig. 37); the synapse inference algorithm was then run on the modified database to construct a mutant CX model. Although descriptions of the *no bridge* mutant suggest that several of the medial glomeruli are not present, our model does not alter any of the other known neurons in CX. The effects of the mutation on the response of the PB projection neuron families can be observed by comparing the mutant model output in Fig. 38 to Fig. 35. As the BU-EB neurons do not receive any input from other neurons in the generated model, their responses in the mutant model (Fig. 39) are identical to those in the original model.

**Fig. 37.**
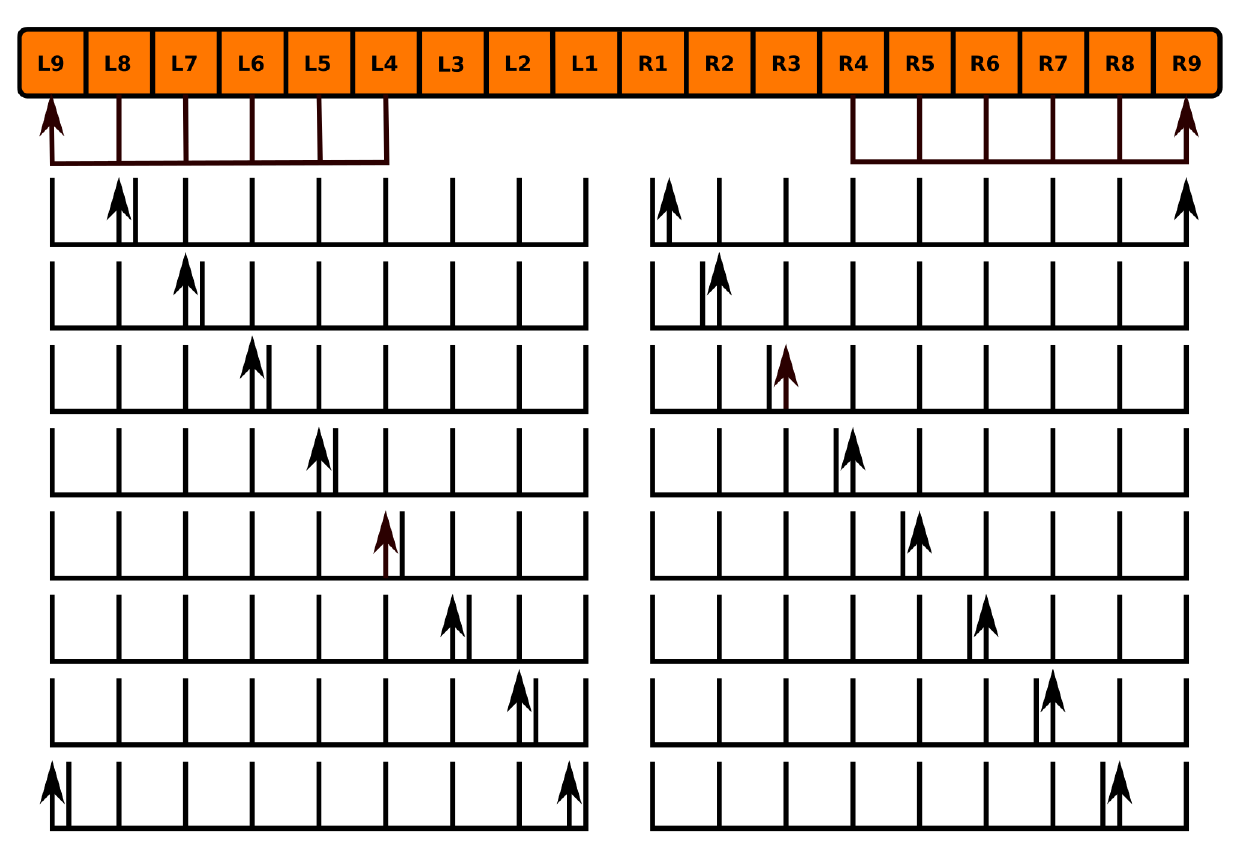
Hypothesized innervation pattern of PB local neurons in no bridge mutant (Table 30).

**Table 30.** Hypothesized PB local neurons in the *no bridge* mutant.

|  | **Label** |
| --- | --- |
| 1 | PB/R9/b-PB/R[4-8]/s |
| 2 | PB/L9/b-PB/L[4-8]/s |
| 3 | PB/L2/b-PB/L[1-9]/s |
| 4 | PB/R7/b-PB/R[1-9]/s |
| 5 | PB/L7/b-PB/L[1-9]/s |
| 6 | PB/R2/b-PB/R[1-9]/s |
| 7 | PB/L3/b-PB/L[1-9]/s |
| 8 | PB/R6/b-PB/R[1-9]/s |
| 9 | PB/L6/b-PB/L[1-9]/s |
| 10 | PB/R3/b-PB/R[1-9]/s |
| 11 | PB/L4/b-PB/L[1-9]/s |
| 12 | PB/R5/b-PB/R[1-9]/s |
| 13 | PB/L5/b-PB/L[1-9]/s |
| 14 | PB/R4/b-PB/R[1-9]/s |
| 15 | PB/R8/b-PB/R[1-9]/s |
| 16 | PB/L1\|L9/b-PB/L[1-9]/s |
| 17 | PB/L8/b-PB/L[1-9]/s |
| 18 | PB/R1\|R9/b-PB/R[1-9]/s |

**Fig. 38.**
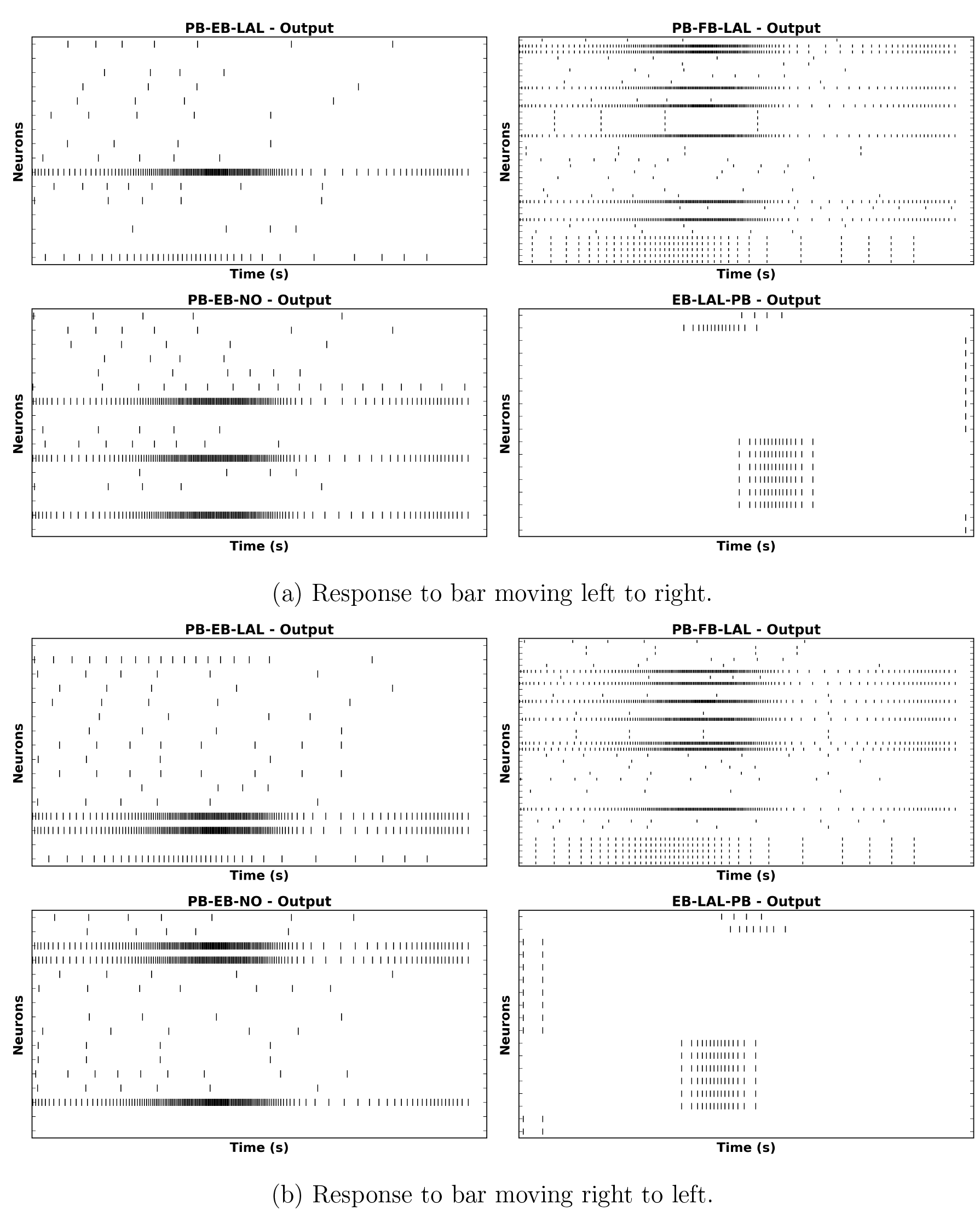
Response of CX projection neurons innervating PB in constructed *no bridge* mutant CX model to moving bar input.

**Fig. 39.**
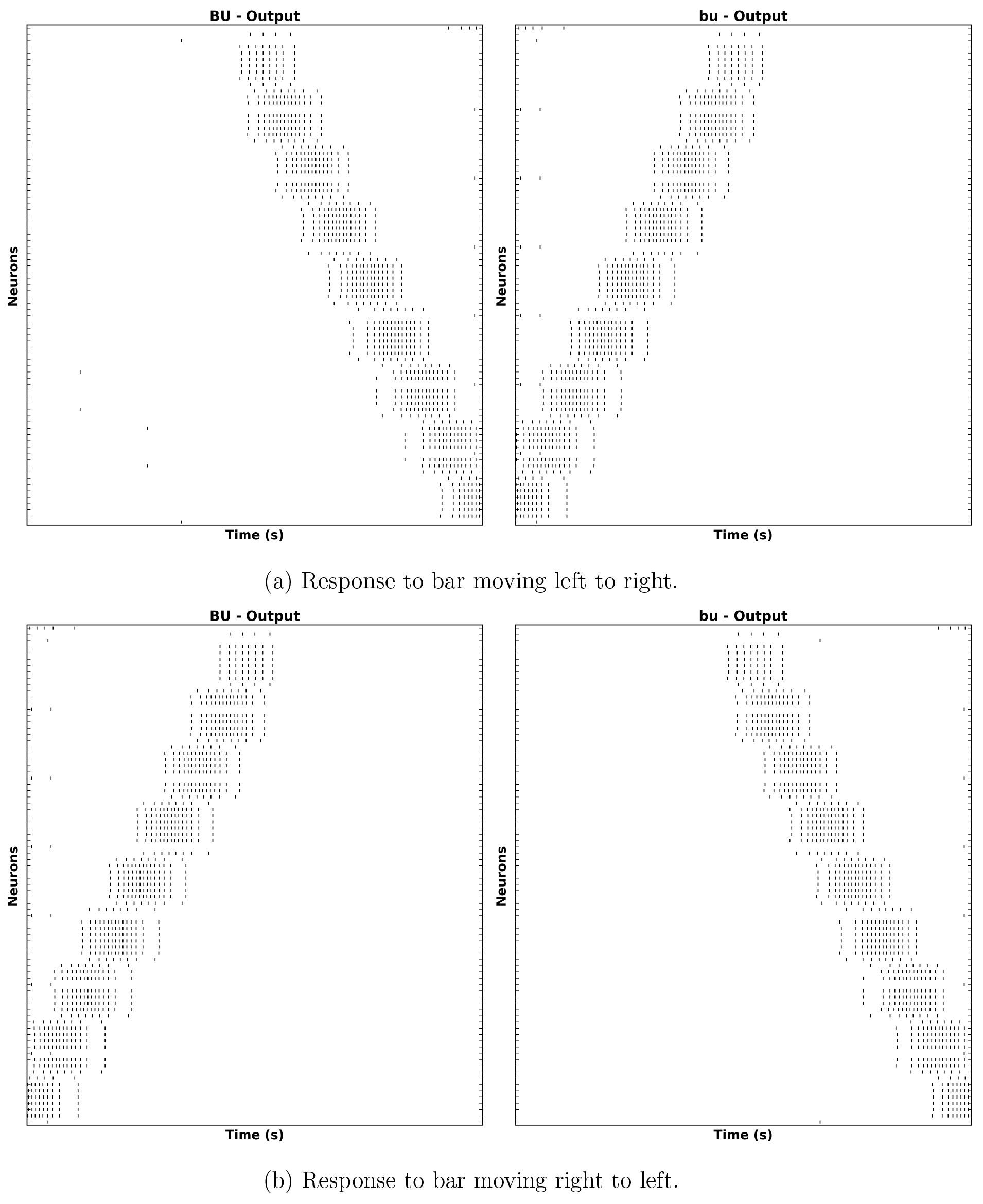
Response of CX projection neurons innervating BU/bu to moving bar input.

### 6.5 Related Work

Since the synapse inference algorithm described above relies entirely on binary arborization overlap information, it cannot infer the number of synapses between neurons. Ongoing work by the developers of the FlyCircuit database [3] that utilizes neuron morphology to infer the number of synapses between neurons will enable the construction of more biologically informed executable circuit models of the central complex [^2^ C.C. Lo, private communication]

## 7 Acknowledgements

The research detailed in this document was supported in part by the AFOSR under grant #FA9550-12-10232, in part by NSF under grant #1544383 and in part by the Department of Electrical Engineering at Columbia University.

